# Dissociating Resting-State Networks of Pedohebephilic Preference and Child Sexual Offending Using Network-Based Prediction

**DOI:** 10.64898/2026.10.21.760530

**Authors:** Emin Serin, Kainat Raza Khan, Tillman H. C. Krueger, Christopher Sinke, Klaus M. Beier, Martin Walter, Boris Schiffer, Till Amelung, Henrik Walter

**Affiliations:** Charité-Universitätsmedizin Berlin, corporate Member of Freie Universität Berlin and Humboldt-Universität zu Berlin, Department of Psychiatry and Neurosciences CCM, Berlin, Germany; German Center for Mental Health (DZPG), Partner Site Berlin - Potsdam, Germany; Freie Universität Berlin, Department of Education and Psychology, Berlin, Germany; Department of Psychiatry, Social Psychiatry and Psychotherapy, Division of Clinical Psychology and Sexual Medicine, Hannover Medical School, Hannover, Germany; Centre of Systems Neuroscience (ZSN), Hannover, Germany; Division of Clinical Psychology and Sexual Medicine, Department of Psychiatry, Social Psychiatry and Psychotherapy, Hannover Medical School, Hannover, Germany; Charité – Universitätsmedizin Berlin, corporate member of Freie Universität Berlin and Humboldt-Universität zu Berlin, Institut für Sexualwissenschaft und Sexualmedizin, Berlin, Germany; Department of Psychiatry and Psychotherapy, Jena University Hospital, Jena, 07743, Germany; Department of Psychiatry, Psychotherapy and Preventive Medicine, Division of Forensic Psychiatry, LWL-University Hospital, Bochum, Germany; Medical School Berlin, Hochschule für Gesundheit und Medizin, Department Psychologie, Germany

**Keywords:** pedophilia, pedohebephilia, child sexual offending, machine learning, functional connectivity, resting-state, forensic psychiatry

## Abstract

Pedohebephilia (P+), the sexual preference for prepubescent and early pubertal minors, regularly causes distress in those affected. While associated with child sexual offending (CSO), P+ is not a prerequisite for CSO. Limited neuroimaging studies have identified alterations associated with P+, while these may primarily reflect offending history. The correlates of P+ independent of CSO therefore remain unclear. We examined this distinction using intrinsic functional networks within a multivariate predictive framework. Using resting-state functional connectivity (rsFC) and network-based prediction (NBS-Predict), we performed pairwise classifications among non-offending pedohebephilic men (P+CSO-, *n*=35), pedohebephilic men with CSO (P+CSO+, *n*=38), and non-pedohebephilic controls without CSO (P-CSO-, *n*=63). Classification-relevant subnetwork nodal strength was associated post hoc with trait impulsivity within P+ groups. P+CSO- were significantly distinguished from P-CSO- (balanced accuracy [BAC]=.621, *p_FDR_*=.041), and showed reductions between default-mode hubs and temporoparietal somatomotor and ventral attention regions. P+CSO+ were also significantly distinguished from P+CSO- (BAC=.645, *p_FDR_*=.032) and yielded a smaller subnetwork centered on the substantia nigra. Against P-CSO-, P+CSO+ showed widespread reductions, particularly across subcortical, default mode, visual, limbic, and somatomotor regions (BAC=.640, *p_FDR_*=.032). Right precentral gyrus nodal strength correlated with impulsivity in P+CSO+ only (*r* = .57, *p_FDR_*=.024). Whole-brain rsFC contains information, distinguishing pedohebephilic preference in the absence of known CSO, while P+CSO+ show additional, more widespread network alterations involving systems implicated in action selection and behavioral regulation. These findings open new avenues for integrating pedohebephilia into broader neurobiological research frameworks.

## INTRODUCTION

Pedohebephilia (P+) refers to a sexual preference for prepubertal and early pubescent minors^1,2^, with reported point prevalence of 1-14% in the general male population^3,4^. Although associated with child sexual offending (CSO), P+ is not a necessary prerequisite^2^. Most individuals with pedohebephilic interests report never acting on them^5^, while only approximately half of individuals convicted of CSO show clinical signs of P+^6^. Distinguishing sexual preference from offending behavior is therefore essential, as neural alterations observed in P+ individuals with a history of CSO (P+CSO+) cannot necessarily be attributed to the sexual preference itself. Studying P+ who have not committed CSO (P+CSO-) provides a critical opportunity to characterize neural features associated with pedohebephilic preference in the absence of known sexual offending.

Previous neuroimaging research has only partly addressed this distinction. Structural alterations involving frontolimbic, temporoparietal, subcortical, and white-matter systems have repeatedly been reported in samples of individuals with pedophilic or pedohebephilic preference^7–14^. However, many of these studies predominantly or exclusively included individuals with a history of CSO. The reviews of direct comparisons between offending and non-offending P+ individuals suggest that some abnormalities previously attributed to pedophilia may instead be more strongly associated with offending history^15,16^. For instance, disrupted white-matter integrity in fronto-occipital, frontotemporal, and callosal pathways distinguishes P+CSO+ from both non-pedohebephilic controls without CSO (P-CSO-) and P+CSO- individuals^12–14^. Similarly, structural abnormalities reported in frontolimbic, temporoparietal, and subcortical regions of P+CSO+ samples^7–11^ have not consistently been reproduced when CSO history is explicitly controlled^11,17^. Together, these findings indicate that the neural correlates of pedohebephilic preference itself remain insufficiently characterized. Moreover, psychiatric comorbidities, including mood, anxiety, and substance-related disorders, occur at elevated rates in pedophilic populations irrespective of offending status^18,19^, and further complicate attempts to isolate neural characteristics associated specifically with sexual preference.

Most neuroimaging studies in this field have focused on brain structure, while comparatively less is known about intrinsic functional network organization. Resting-state functional magnetic resonance imaging (rs-fMRI) captures spontaneous brain activity without explicit tasks, is temporally reliable^20^, overlaps substantially with task-evoked functional systems^21^, and has been widely used to characterize distributed alterations associated with neuropsychiatric conditions^22–24^. Only three resting-state studies have investigated pedophilic or pedohebephilic populations^25–27^. Kärgel et al.^25^ used predefined seed regions and reported reduced connectivity of the posterior cingulate cortex and left amygdala in P+CSO+ relative to both P+CSO- and P-CSO- participants, and concluded that these alterations relate to CSO rather than to P+ itself. However, the seed-based design may have missed network-level alterations. Complementing this, Cantor et al.^26^ subsequently applied independent whole-brain component analysis (ICA) and reported altered connectivity in the default mode (DMN), frontoparietal (FPN), insular, and occipital regions. However, the P+ sample in this study entirely consisted of individuals with a history of CSO, compared against non-sexual offenders and individuals without criminal history. While this design could control for general criminality, it could not separate sexual preference from offending due to the lack of P+CSO-. Another study by Andersson et al.^27^ also applied ICA and reported widespread reductions in visual, sensorimotor, auditory, and cerebellar networks in a mixed sample of P+, which does not allow for disentangling the effects of P+ from CSO. Moreover, all studies were limited by relatively small P+ samples. Consequently, whole-brain resting-state network organization associated with pedohebephilic preference in individuals without CSO remains largely unexplored.

Disentangling these distinctions additionally requires an analytic framework suited to distributed signals. Higher-order functions and psychiatric conditions are encoded across networks of multiple regions rather than isolated regions^28–31^. Conventional mass-univariate approaches, such as general linear models (GLM), are well suited to identifying localized group differences yet may not capture multivariate patterns distributed across the functional network^32,33^. Multivariate machine learning (ML) approaches, in contrast, integrate high-dimensional brain data into a single predictive model and complement GLM in two ways: out-of-sample evaluation to estimate generalizability of findings, and prediction at the level of individuals^32,34^. To date, only three reports have applied machine learning to the prediction of P+, using task-based fMRI^35,36^ or diffusion MRI^37^, and none used resting-state networks. The two fMRI studies^35,36^ analyzed the same 56 participants from a single site and reported high accuracies for P+ itself (93–95%). Their discriminative patterns involved distributed cingulate, temporal, parietal, occipital, and basal-ganglia regions, several of which also contributed to the present classifications. However, the 24 P+ participants comprised both offending and non-offending individuals; therefore, these patterns do not establish a pedophilia-specific neural signature independent of offending. The diffusion MRI study^37^ distinguished 14 P+CSO+ participants from 15 non-pedophilic controls with a balanced accuracy of 75.5% and identified predictive features involving anterior cingulate and prefrontal–amygdala pathways; nevertheless, the resulting white-matter pattern could not separate the effects of pedophilia from those of offending. Notably, the symptom severity of comorbid psychiatric conditions, including anxiety and depression, was not controlled in the previous studies, which may increase the likelihood that classifications were driven by group differences in such psychiatric symptoms.

The present study addresses these limitations by using resting-state functional connectivity and network-based predictive modeling, NBS-Predict^38^, in a comparatively large, multisite sample of P+CSO-, P+CSO+, and P-CSO- participants (Fig. 1). Using NBS-Predict, we classified all three pairwise groups. This design isolates the network signature of pedohebephilic preference from that of CSO: the contrast against P-CSO- in the absence of offending identifies the pedohebephilic preference; contrast against P+CSO+ identifies additional differences associated with CSO history among P+ individuals; and P+CSO+ vs. P-CSO- provides complementary characterization of the combined P+ and CSO phenotype. Given the heterogeneity of prior findings, we adopted a data-driven, exploratory approach without a priori regional hypotheses. As a secondary, post hoc question, we examined whether the identified subnetworks relate to trait impulsivity within the P+ groups, which has been repeatedly shown to be altered, particularly among CSO individuals^39–41^.

**Figure 1.**
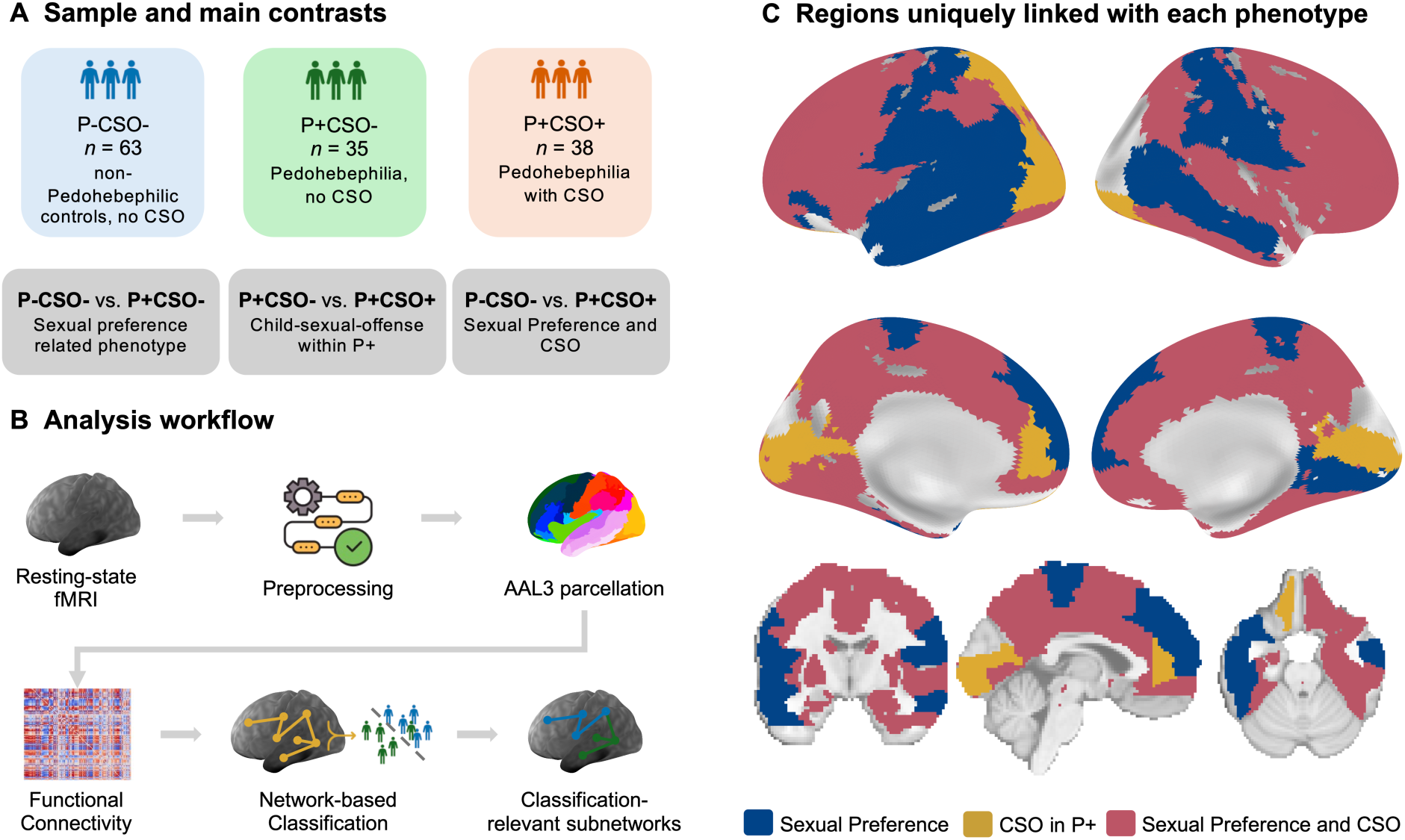
Study design, analysis workflow, and regions uniquely associated with each phenotype. **A**. Sample composition and the pairwise classification contrasts. Non-pedohebephilic men without a history of child sexual offending (P−CSO−, *n*=63), men with pedohebephilia without CSO (P+CSO−, *n*=35), and men with pedohebephilia with CSO (P+CSO+, *n*=38). Each contrast isolates a different phenotype: P−CSO− vs. P+CSO− pedohebephilic sexual preference in the absence of documented contact offending, P+CSO− vs. P+CSO+ child sexual offending within pedohebephilia, and P−CSO− vs. P+CSO+ the combined phenotype. **B.** Analysis workflow. Resting-state fMRI data were preprocessed with ENIGMA HALFpipe and parcellated using the AAL3 atlas without the cerebellum. The resulting functional connectivity matrices were entered into NBS-Predict, which performs network-based feature selection and classification within nested cross-validation, yielding weighted, classification-relevant subnetwork per contrast. **C.** Regions selected in the classification-relevant subnetwork of one contrast only, displayed on the surface and in volumetric space. Blue, sexual interest (P−CSO− vs. P+CSO−); yellow, CSO within pedohebephilia (P+CSO− vs. P+CSO+); red, combined phenotype (P−CSO− vs. P+CSO+). Grey indicates regions that were either not selected in any contrast or selected in more than one, and are therefore not phenotype-specific.

## METHODS

### Subjects

Male participants were recruited through the multisite Neural Mechanisms Underlying Pedophilia and Sexual Offending against Children (NeMUP) collaboration across four German sites: Berlin, Hannover, Duisburg-Essen, and Kiel. Participants were recruited through the Prevention Project Dunkelfeld, prisons, online forums, advertisements, and mailing lists. In this study, both pedophilia and hebephilia, which are defined as sexual preference in prepubescent and early-pubescent bodies, were used; we refer to them as pedohebephilic (P+) throughout^i^. Pedohebephilia was diagnosed according to ICD-10 criteria^42^, whose age criterion for preferred partners extends into early puberty, and both constructs often converge in several domains, including sexual behavior and arousal^43^. CSO was defined as at least one physical sexual offense against a child under 14 years, verified by self-report or legal records. General cognitive ability was assessed using the Wechsler Adult Intelligence Scale-IV^44^. Comorbid mental disorders were assessed with the Structured Clinical Interview for DSM-IV-TR (SCID-I/II)^45,46^ and the Hamilton Anxiety (HAMA)^47^ and Depression (HAMD)^48^ Scales. Exclusion criteria comprised an acute mental disorder diagnosed using the SCID, severe anxiety (HAMA ≥ 25) or depression (HAMD ≥ 15), brain injury, neurological illness, diagnostic uncertainty regarding pedophilia, current psychotropic medication including androgen-deprivation therapy, and poor MRI quality (Supplementary Methods). The initial sample comprised 76 P-CSO-, 50 P+CSO+, and 51 P+CSO- participants; after exclusions, the final sample included 63 P-CSO-, 38 P+CSO+, and 35 P+CSO- participants (Fig. 1a). 20 of the 63 P-CSO- participants were previously used by Kneer et al.^49^, comparing P+CSO- with P-CSO- using seed-based (i.e., amygdala) resting-state functional connectivity analysis. None of the P+ participants in the present study were included in that report.

### MRI Image Acquisition and Processing

Resting-state MRI scans were acquired on 3T MRI scanners across four sites using a standardized protocol. Anatomical images were obtained using a T1-weighted magnetization-prepared rapid gradient-echo (MP-RAGE) sequence with an isotropic spatial resolution of 1mm³ (TR/TE=2500/4.33ms, flip angle=7°). A total of 275 resting-state volumes were acquired using an echo-planar imaging (EPI) sequence with a spatial resolution of 2.3 × 2.3 × 3.3mm³ (TR/TE=2400/30ms, flip angle=80°). Preprocessing was performed using ENIGMA HALFpipe (version 1.2.2)^50^, a standardized pipeline integrating FSL, FreeSurfer, and fMRIPrep. Specifically, the preprocessing steps involved: skull stripping using FreeSurfer; slice timing and motion correction; normalization using ANTs^51^; grand mean scaling; further noise removal using ICA-AROMA (non-aggressive)^52^, temporal filtering using Gaussian high-pass filters (125 s) and smoothing (4mm FWHM kernel). Discrete cosine filters (i.e., temporal filtering) and noise components detected by ICA-AROMA were regressed out simultaneously in a single regression model to prevent spurious artefacts^53^. The quality of preprocessed images was assessed through visual inspection and objective evaluation of preprocessing steps, and participants with inadequate quality control ratings or frame-wise displacement (FD) rate^54^ higher than .50 were excluded from further analyses. Following preprocessing, resting-state time series of each brain region were extracted using the Automated Anatomical Labeling Atlas (AAL3v2)^55^. Due to the cerebellar cutoff in the brain images, the cerebellum was excluded from the analyses, leaving a total of 140 regions. Pairwise correlations between regional time series were computed to create a 140×140 rsFC matrix (9,730 connections) for each subject and were normalized using Fisher’s *r*-to-*z* transform, subsequently used as the feature set for classifications. Detailed acquisition and preprocessing parameters are provided in the Supplementary Methods.

### Network-based Prediction

To distinguish each group pair (P-CSO- vs. P+CSO-, P-CSO- vs. P+CSO+, P+CSO- vs. P+CSO+) and identify key regions and connections contributing to classification, we employed NBS-Predict (v1.0.0-beta.13), a network-based predictive modeling approach integrating graph theory with ML^38^. NBS-Predict uses resting-state functional connectivity (rsFC) matrices as input, selects subnetworks associated with group differences, and applies ML algorithms within nested cross-validation (CV). As an extension of the network-based statistic (NBS)^56^, which provides greater statistical power than mass-univariate testing^57^, NBS-Predict incorporates NBS into feature selection. This leverages GLM with group-level contrasts (e.g., P-CSO->P+CSO+) to identify subnetworks associated with group differences, which enables straightforward interpretation. The directional contrasts corresponding to the identified networks with reduced connections were P-CSO->P+CSO-, P+CSO->P+CSO+, and P-CSO->P+CSO+. To validate primary model contrasts, three models with reversed contrasts (e.g., P-CSO-<P+CSO+) were trained. The feature selection threshold was *p*<.01 to retain the most robust connections. Potential confounding effects of age, site, HAMA, and HAMD scores were controlled using cross-validated confound regression^58^ (Table 1).

**Table 1.**
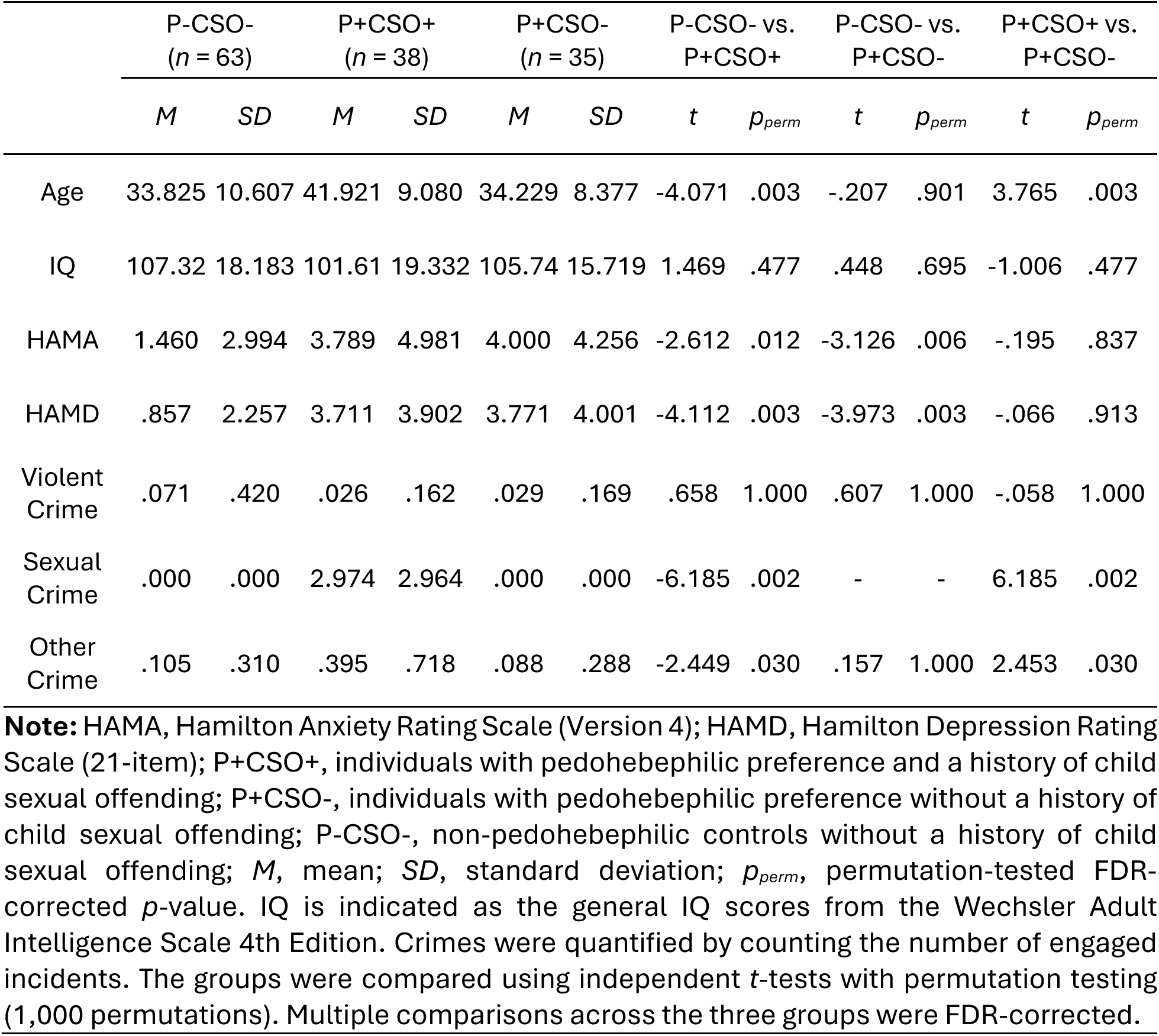
Demographics, clinical, and offense-related characteristics across groups.

|  | P-CSO-<br>(*n* = 63) |  | P+CSO+<br>(*n* = 38) |  | P+CSO-<br>(*n* = 35) |  | P-CSO- vs.<br>P+CSO+ |  | P-CSO- vs.<br>P+CSO- |  | P+CSO+ vs.<br>P+CSO- |  |
|---|---|---|---|---|---|---|---|---|---|---|---|---|
|  | *M* | *SD* | *M* | *SD* | *M* | *SD* | *t* | *p<sub>perm</sub>* | *t* | *p<sub>perm</sub>* | *t* | *p<sub>perm</sub>* |
| Age | 33.825 | 10.607 | 41.921 | 9.080 | 34.229 | 8.377 | -4.071 | .003 | -.207 | .901 | 3.765 | .003 |
| IQ | 107.32 | 18.183 | 101.61 | 19.332 | 105.74 | 15.719 | 1.469 | .477 | .448 | .695 | -1.006 | .477 |
| HAMA | 1.460 | 2.994 | 3.789 | 4.981 | 4.000 | 4.256 | -2.612 | .012 | -3.126 | .006 | -.195 | .837 |
| HAMD | .857 | 2.257 | 3.711 | 3.902 | 3.771 | 4.001 | -4.112 | .003 | -3.973 | .003 | -.066 | .913 |
| Violent Crime | .071 | .420 | .026 | .162 | .029 | .169 | .658 | 1.000 | .607 | 1.000 | -.058 | 1.000 |
| Sexual Crime | .000 | .000 | 2.974 | 2.964 | .000 | .000 | -6.185 | .002 | - | - | 6.185 | .002 |
| Other Crime | .105 | .310 | .395 | .718 | .088 | .288 | -2.449 | .030 | .157 | 1.000 | 2.453 | .030 |

**Note:** HAMA, Hamilton Anxiety Rating Scale (Version 4); HAMD, Hamilton Depression Rating Scale (21-item); P+CSO+, individuals with pedohebephilic preference and a history of child sexual offending; P+CSO-, individuals with pedohebephilic preference without a history of child sexual offending; P-CSO-, non-pedohebephilic controls without a history of child sexual offending; *M*, mean; *SD*, standard deviation; *p<sub>perm</sub>*, permutation-tested FDR-corrected *p*-value. IQ is indicated as the general IQ scores from the Wechsler Adult Intelligence Scale 4th Edition. Crimes were quantified by counting the number of engaged incidents. The groups were compared using independent *t*-tests with permutation testing (1,000 permutations). Multiple comparisons across the three groups were FDR-corrected.

Three linear classification algorithms were evaluated: support vector classification (SVC), logistic regression (LR), and linear discriminant analysis (LDA). Predictive models were evaluated using 50 repeated 7×7 nested cross-validation, with balanced accuracy (BAC) as the primary metric, along with complementary AUC-ROC and F1 metrics. Statistical significance of classification performance was determined via permutation testing (1,000 permutations)^59^ and FDR-corrected across ML algorithms and classification pairs^60^. Following classification, NBS-Predict identified the subnetworks most relevant to each class, producing a weighted network (values 0-1) that reflects each connection’s contribution to the classification. For visualization only, a weight threshold of .97 was applied to retain connections with consistently high classification involvement; it did not affect classifier performance. Detailed NBS-Predict methodology is provided elsewhere^61^. Nodal strength (NS), analog of nodal degree in weighted networks, was calculated to quantify each region’s centrality and connectivity within the identified networks (Supplementary Methods). Anatomical regions from the AAL3 atlas^55^ were additionally mapped onto canonical resting-state functional networks^62^ to facilitate network-level interpretation (Supplementary Table 1 for region-network mapping). To assess whether the classification findings depended on the choice of brain parcellation^63^, the complete ML analyses were repeated using the Brainnetome atlas^64^ (Supplementary Figures 1-3, Supplementary Table 2 for findings). As a post hoc analysis, we investigated the behavioral relevance of the identified functional connectivity patterns by correlating trait impulsivity scores from the Barratt Impulsiveness Scale (BIS-11; total score; Supplementary Methods)^65^ and regional nodal strength separately for P+CSO- and P+CSO+ participants, controlling for acquisition site (FDR corrected across regions).

## RESULTS

### Sample Characteristics

Demographic and clinical characteristics are summarized in Table 1. P+CSO+ participants were significantly older than both other groups and had a greater number of sexual and non-sexual offenses. Both P+ groups had significantly higher HAMA and HAMD scores than P-CSO- participants, with no significant difference between them. Groups did not differ in IǪ or number of violent offenses, and P+CSO- did not differ from P-CSO- in age or offense counts.

### Group Classifications

In all three classifications (P-CSO->P+CSO-, P-CSO->P+CSO+, P+CSO->P+CSO+), at least one classifier significantly distinguished the groups, and SVC performed best on BAC throughout. Consistency across algorithms varied: all three algorithms survived FDR correction for P-CSO->P+CSO+, two of three for P+CSO->P+CSO+, and only SVC for P-CSO- >P+CSO- (Table 2). All reversed-contrast models (e.g., P-CSO- < P+CSO+) failed because no subnetwork survived feature selection, and, therefore, the model could not be trained. Results were largely consistent using the Brainnetome atlas, with discrepancies attributable to differences in the number and definition of delineated regions (Supplementary Figures 1–3, Supplementary Table 2). For instance, AAL3 includes a dedicated substantia nigra region, which Brainnetome does not explicitly define.

**Table 2.** Results of all three pairwise classifications using NBS-Predict.

|  | Method | Balanced Accuracy | *p<sub>FDR</sub>* | AUC-ROC | F1 |
|---|---|---|---|---|---|
| P-CSO- > P+CSO- |  |  |  |  |  |
|  | **SVC** | **.621** | **.041** | **.589** | **.609** |
|  | Logistic Regression | .569 | .096 | .566 | .668 |
|  | LDA | .589 | .063 | .573 | .628 |
| P+CSO- > P+CSO+ |  |  |  |  |  |
|  | **SVC** | **.645** | **.032** | **.600** | **.579** |
|  | **Logistic Regression** | **.623** | **.041** | **.612** | **.542** |
|  | LDA | .612 | .059 | .658 | .618 |
| P-CSO- > P+CSO+ |  |  |  |  |  |
|  | **SVC** | **.640** | **.032** | **.646** | **.748** |
|  | **Logistic Regression** | **.580** | **.041** | **.597** | **.720** |
|  | **LDA** | **.609** | **.041** | **.607** | **.693** |

**Note:** P+CSO+, individuals with pedohebephilic interests and a history of child sexual offending; P+CSO-, individuals with pedohebephilic interests without a history of child sexual offending; P-CSO-, non-pedohebephilic controls without a history of child sexual offending, SVC, support vector classifier; LDA, linear discriminant analysis; AUC-ROC, area under the receiver operating characteristic curve. A 50 repeated 7x7 nested cross-validation was used for model evaluation and hyperparameter optimization. Age, site, HAMD, and HAMA scores were controlled for. P-values were computed using permutation testing (1,000 permutations). FDR correction was applied across multiple classifications and classifiers (9 in total for subgroup classification). The significant classifications (*p* < 0.05) are highlighted. Classifications using reverse subnetworks (e.g., P-CSO- < P+CSO+) failed as no subnetworks survived feature selection, and, thus, the model could not be trained.

### Pedohebephilic vs. non-pedohebephilic non-offenders (P-CSO->P+CSO-)

NBS-Predict using SVC significantly distinguished P+CSO- from P-CSO- participants, achieving a BAC of .621 (*p_FDR_*=.041; Table 2). The model revealed a circumscribed subnetwork of 25 decreased connections among 20 regions in P+CSO- compared to P-CSO-, primarily involving temporal and parietal regions with some frontal involvement (Fig. 2, Supplementary Tables 3, 4). Key connectivity hubs with the highest nodal strength included left (NS=9.904) and right (NS=3.969) middle temporal gyrus (MTG), left (NS=6.919) and right (NS=2.948) supramarginal gyrus (SMG), left rolandic operculum (NS=3.983), and left angular gyrus (NS=3.959). At the network level, relevant regions were primarily involved in the default mode network (DMN), somatomotor network (SMN), and ventral attention (or salience) network (VAN), as well as partially in the visual network (VIS). Regions with high nodal strength were predominantly located in the left hemisphere. No significant associations were observed between regional nodal strength within this subnetwork and BIS-11 total scores in P+CSO- participants (Supplementary Table 5).

**Figure 2.**
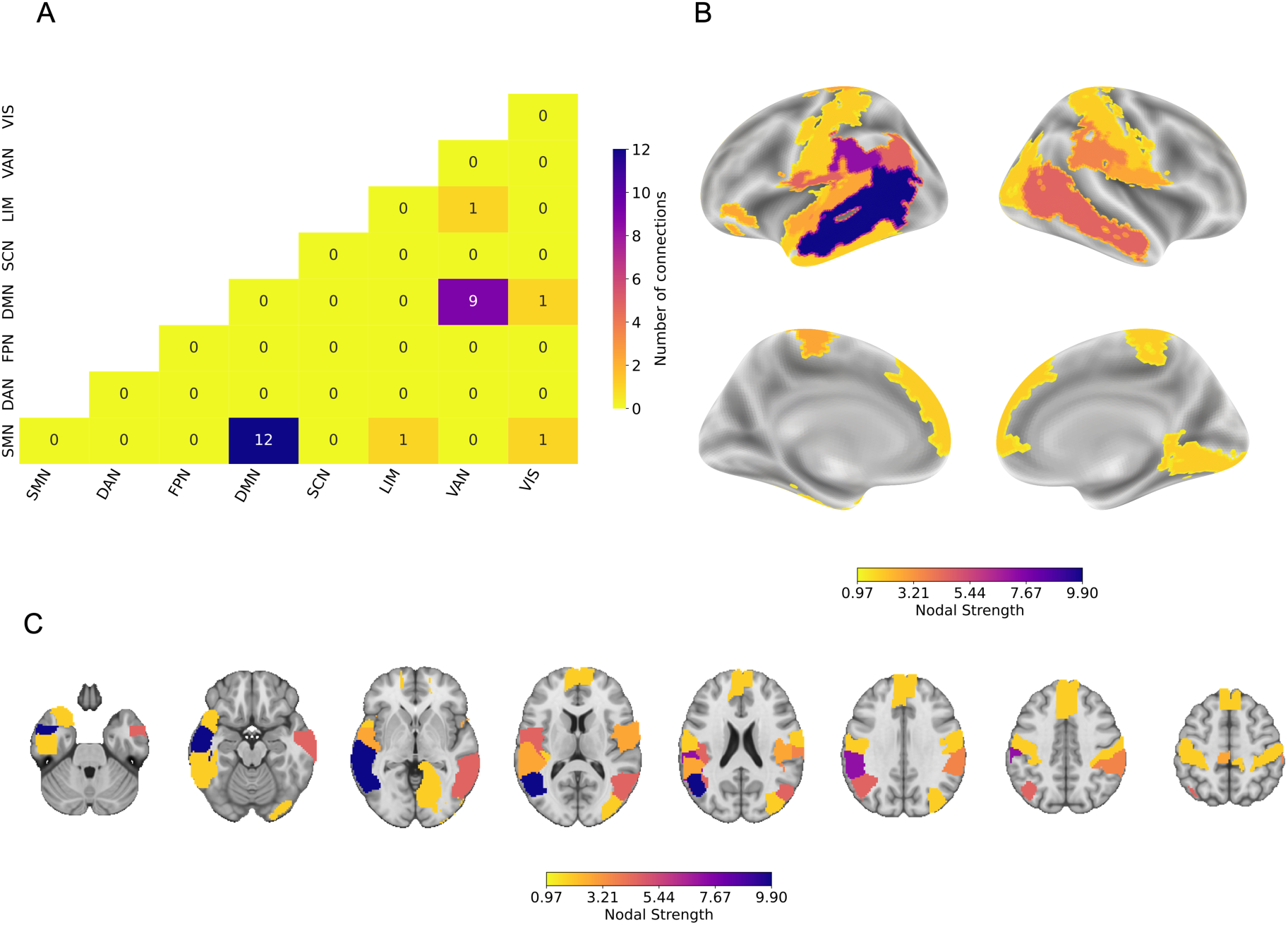
Subnetwork of relevant decreased connections in P+CSO- (25 connections among 20 regions), classifying P+CSO- and P-CSO- groups using a linear support vector classifier in NBS-Predict. A weight threshold of.97 was applied to visualize highly relevant connections for the classification. **A.** The heatmap illustrates the number of connections between canonical brain networks. The region-network mapping is given in Supplementary Table 1. **B.** and **C.** Nodal strength of brain regions in the subnetwork is displayed on the brain surface (fsaverage) and in volumetric space. The details of the nodal strength calculation are provided in the Supplementary Methods.

### Pedohebephilic non-offenders vs. offenders (P+CSO->P+CSO+)

Significant classification between P+CSO- and P+CSO+ was achieved, yielding a BAC of .645 (*p_FDR_*=.032; Table 2). The model identified a small subnetwork comprising 8 connections among 9 regions with reduced functional connectivity in P+CSO+ than in P+CSO- participants (Fig. 3, Supplementary Tables 6, 7). The subnetwork comprised regions involved in visual, default mode, subcortical (SCN), and dorsal attention (DAN) networks. Within the subnetwork, the left substantia nigra pars reticulata (SNpr; NS=5.898) was the subnetwork hub and connected to the left medial OFC, left calcarine gyrus, left superior parietal gyrus, and bilateral middle and right inferior occipital regions. No significant associations between NS within this subnetwork and BIS-11 total scores were observed in P+ (Supplementary Table 8).

**Figure 3.**
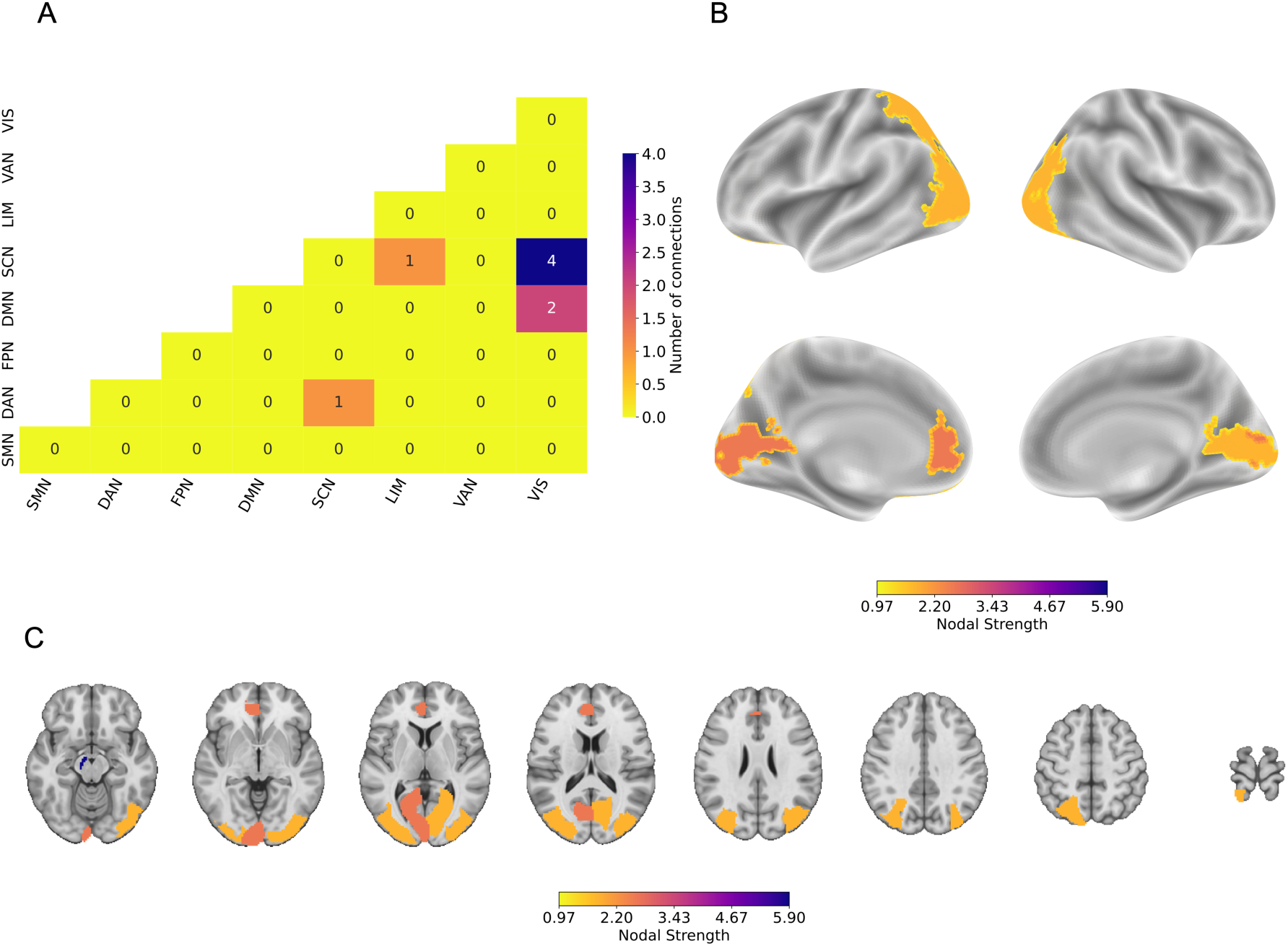
Subnetwork of relevant decreased connections in P+CSO+ (8 connections among 9 regions), classifying P+CSO- and P+CSO+ groups using a linear support vector classifier in NBS-Predict. A weight threshold of .97 was applied to visualize highly relevant connections for the classification. **A.** The heatmap illustrates the number of connections between canonical brain networks. The region-network mapping is given in Supplementary Table 1. **B.** and **C.** Nodal strength of brain regions in the subnetwork is displayed on the brain surface (fsaverage) and in volumetric space. The details of the nodal strength calculation are provided in the Supplementary Methods.

### Pedohebephilic offenders vs. non-pedohebephilic non-offenders (P-CSO->P+CSO+)

SVC also significantly distinguished P+CSO+ from P-CSO- participants, achieving a BAC of .640 (*p*_FDR_=.032; Table 2). This classification revealed a substantially more extensive subnetwork comprising 167 connections with lower functional connectivity in P+CSO+ participants among 89 regions (Fig. 4, Supplementary Tables 9, 10). Connectivity reductions were distributed throughout the whole brain, primarily involving regions from the SCN, DMN, VIS, limbic network (LIM), and SMN. The regions with the highest nodal strength were the left pallidum (NS=30.634), left putamen (NS=20.693), right calcarine gyrus (NS=13.903), left pregenual ACC (NS=11.951), left supracallosal ACC (NS=9.904), right lingual gyrus (NS=9.901), and right anterior OFC (NS=9.861), indicating their role as connectivity hubs. Within P+CSO+ participants, nodal strength of the right precentral gyrus within this subnetwork was positively associated with BIS-11 total impulsivity scores (*r*=.57, *p*_FDR_=.024; Supplementary Table 11, Supplementary Figure 4), which was primarily connected with the right lateral OFC, left putamen, left pallidum, and right nucleus accumbens (NAc).

**Figure 4.**
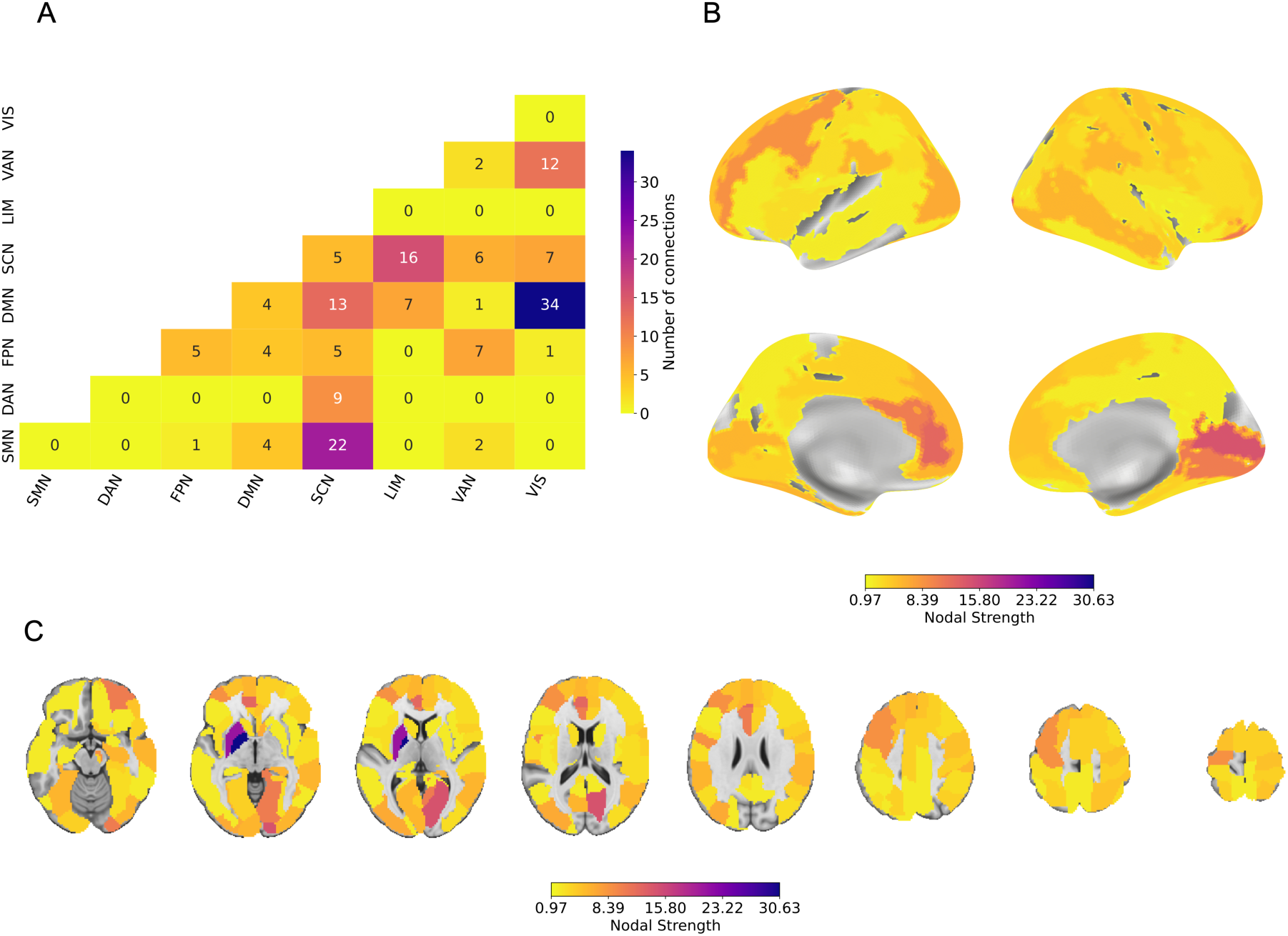
Subnetwork of relevant decreased connections in P+CSO+ (167 connections among 89 regions), classifying P+CSO+ and P-CSO- groups using a linear support vector classifier in NBS-Predict. A weight threshold of.97 was applied to visualize highly relevant connections for the classification. **A.** The heatmap illustrates the number of connections between canonical brain networks. The region-network mapping is given in Supplementary Table 1. **B.** and **C.** Nodal strength of brain regions in the subnetwork is displayed on the brain surface (fsaverage) and in volumetric space. The details of the nodal strength calculation are provided in the Supplementary Methods.

## DISCUSSION

Using resting-state functional connectivity and network-based predictive modeling, we distinguished pedohebephilic men with and without a history of child sexual offending from non-pedohebephilic controls and from one another (Fig. 1c). Our main finding was that P+CSO- could be distinguished above chance from P-CSO-, indicating a pedohebephilic-specific signature. Specifically, pedohebephilic preference was associated with a circumscribed subnetwork linking default mode hubs to temporoparietal somatomotor and ventral attention regions. The two additional contrasts complemented this finding by identifying further network alterations associated with CSO history among P+ individuals. These alterations were largely distinct from the P+CSO- pattern and substantially involved subcortical, somatomotor, attentional, and frontoparietal systems. Together, the findings suggest that network characteristics associated with pedohebephilic preference can be detected in the absence of known sexual offending, while offending history within pedohebephilia is associated with additional alterations in distributed functional networks.

### Network Characteristics Associated with Pedohebephilic Preference in the Absence of CSO

The P+CSO- vs. P-CSO- contrast constitutes the principal classification of the present study as it examines network characteristics associated with pedohebephilic preference without the additional presence of CSO. Above-chance classification yielded a network of reduced connectivity between default mode hubs (e.g., middle temporal and angular gyri), somatosensory, and opercular regions. These regions are also present in the P+CSO+ subnetwork (Supplementary Table 9), consistent with their relevance to pedohebephilic preference irrespective of offending history. These findings also converge with structural evidence. Abé et al.^10^ reported reduced surface area in bilateral medial superior frontal, postcentral, and supramarginal gyri, and the left inferior frontal gyrus, in an independent sample of men with pedophilic disorder comprising both offending and non-offending individuals.

Network alterations reported in other neuropsychiatric conditions also caution against inferring functional specificity. For instance, altered DMN–somatosensory connectivity has been observed across multiple clinical domains, including schizophrenia and depression^23,66,67^. These alterations were independent of depression severity in our study, as HAMD scores were regressed out, and in previous research^68,69^, making comorbid depression an unlikely explanation and instead suggesting potential transdiagnostic characteristics. Notably, DMN-somatosensory connectivity has been shown to have high predictive relevance for autism spectrum disorder (ASD)^70^. Empirical and clinical evidence links ASD and pedohebephilic disorder^71,72^, and these two conditions share features that likely map onto this DMN-somatosensory connectivity: social interaction difficulties, generalized in ASD, while adult-focused in pedohebephilia^73,74^, and highly idiosyncratic interests^6^, reminiscent of the special interests characteristic of ASD, whereas focused on sexual arousal in P+.

Two further observations connect this network to sexuality particularly. In both P+ groups against P-CSO-, altered subnetworks involved the opercular cortex, which is implicated in sexual arousal, sexual auras in focal epilepsy, and genital somatosensory processing^75–78^. Given the involvement of the DMN in self-referential processing^79^, reduced coupling between default mode hubs and these regions might relate to atypical self-perception; research indicates a more child-like self-concept in many individuals with pedohebephilia that may expand to imagined sexual scenarios^80,81^, which aligns with clinical explanations of individuals imagining themselves as children within sexual scenarios. Therefore, this subnetwork may support the embedding of bodily and arousal-related signals into self-related representations, and its alteration may contribute to the atypical social functioning, arousal patterns, and self-perception that characterize P+.

These interpretations nonetheless require considerable caution. This contrast yielded the weakest model in the study (BAC=0.621) and was the only contrast in which just one algorithm survived FDR correction. While it supports the claim that a pedohebephilia-related connectivity signature is detectable above chance in a whole-brain, data-driven analysis with offending history held constant, it could neither provide inference about any individual nor any clinical or forensic application.

### Additional Network Alterations Associated with CSO in Pedohebephilia

The classification between P+ groups complements the principal findings of P+ associated functional networks, as these groups share pedohebephilic preference but differ in CSO history. The classification identified a small subnetwork centered on SNpr (substantia nigra pars reticulata), which showed reduced connectivity in P+CSO+ with prefrontal, limbic, parietal, and occipital regions. The SNpr is a major output structure of the basal ganglia and has a well-established role in the regulation of action selection and movement through inhibitory projections via thalamic nuclei^82^. Recent research has reliably established its connectivity with frontal, parietal, and occipital regions^83^, mirroring the patterns identified in the present study. The present findings therefore may suggest that impaired cognitive modulation of action selection and suppression differentiates offending from non-offending P+ individuals.

This interpretation is further supported by the broader P+CSO+ vs. P-CSO- classification. In contrast to the relatively circumscribed network associated with P+CSO-, P+CSO+ showed widespread reductions encompassing 167 connections across cortical, limbic, and subcortical regions. Regions with the greatest nodal strength were localized to subcortical regions (i.e., left pallidum and putamen) together with anterior cingulate, orbitofrontal, visual, and somatomotor regions. The abnormalities within frontostriatal and related networks, which contribute to action selection and cognitive control, have been previously reported in neuroimaging studies of P+CSO+^7–9,16,84^. Supporting this, previous behavioral studies from the NeMUP cohort and other samples have demonstrated poorer inhibitory control in offending than non-offending P+ individuals^40,41,85–87^. Consistently, only in P+CSO+ (when contrasted against P-CSO-), nodal strength within the identified subnetwork correlated with trait impulsivity, and only for the right precentral gyrus, which connected primarily to the pallidum, NAc, and OFC. Within this subnetwork of reduced connectivity compared to P-CSO-, the positive association between precentral nodal strength and impulsivity scores, measured with BIS-11, may reflect a shortcut biasing network dynamics toward motor readiness at the expense of inhibitory signaling. Importantly, this finding should nonetheless be considered preliminary, as the analysis was performed post hoc with 38 P+CSO+ participants and, therefore, requires further investigation in larger samples.

Collectively, the two contrasts involving P+CSO+ suggest additional alterations beyond the network characteristics observed in P+CSO- participants, particularly in subcortical and cortical systems involved in action selection and behavioral regulation. Nevertheless, these findings should be interpreted as associations with CSO history rather than causal neural mechanisms, given the cross-sectional design of this study.

### Classification Performance

The accuracies reported here are lower than those in previous ML studies of pedophilia using diffusion imaging or task-evoked brain activations^35–37^. However, the present sample is larger than those of earlier studies (*n*=136 vs. *n*=29-56). Small neuroimaging datasets can produce unstable and upwardly biased estimates of classification accuracy, especially when model selection and validation are not sufficiently separated^34,88,89^. The more conservative estimates observed here may therefore partly reflect reduced sampling variability and provide a more realistic estimate of out-of-sample performance. Additionally, P+ co-occurs frequently with psychiatric comorbidity, particularly mood and anxiety disorders, irrespective of offending status^18,19^, which was supported by the elevated HAMA and HAMD scores in both P+ groups in this study. Accordingly, unlike the earlier studies, which covaried only age and intelligence^35–37^, HAMA and HAMD scores together with age and acquisition site were regressed out, removing shared variance that earlier classifiers could have exploited. Nevertheless, remaining confounding by unmeasured clinical, developmental, or behavioral characteristics cannot be excluded. Moreover, direct comparison with previous studies remains limited by substantial methodological differences, including imaging modality, feature dimensionality, sample composition, diagnostic definitions, and validation procedures. Accordingly, lower classification performance should not be taken as straightforward evidence of less pronounced underlying neurobiological differences.

### Limitations and Future Directions

Several methodological limitations should be considered when interpreting the present findings. While the present sample is the largest to date in this literature, it remains modest for ML applications, particularly given the dimensionality of whole-brain functional connectivity. The significant yet modest classification performance and absence of an independent validation cohort limit diagnostic and forensic utility. In addition, the present study used a single neuroimaging modality. Pedohebephilic preference and offending behavior may be unlikely to be captured comprehensively by a single modality alone, and, therefore, integrating multi-modal measures could provide comprehensive characterization of the relevant neurobiology^90,91^. The sample comprises both pedophilic and hebephilic preferences, which may increase within-group heterogeneity and limit comparability with previous studies that focused exclusively on pedophilia. Given this heterogeneity in clinical presentation, together with comorbidity and neurobiology, frameworks that model inter-individual variability explicitly, such as normative modeling^92^, may be better suited to this population. Moreover, P+CSO- participants were recruited substantially through a prevention program and are predominantly help-seeking, which may not generalize to P+CSO- who do not seek treatment. Furthermore, while the present three-group design allows us to distinguish findings associated with sexual preference from offending status, it cannot estimate a general effect of CSO regardless of P+. To this end, future studies should include a P-CSO+ sample of comparable size.

## CONCLUSIONS

Using whole-brain resting-state functional connectivity and network-based machine learning, we dissociated pedohebephilic preference from child sexual offending in out-of-sample classification. The two phenotypes were associated with largely distinct sets of connections: pedohebephilic preference with a circumscribed network linking default-mode hubs to temporoparietal somatomotor and ventral attention regions, and offending history within pedohebephilia with subcortical systems centered on the substantia nigra. The combined phenotype against controls showed widespread reductions, with strongest hubs in the pallidum and putamen. Trait impulsivity was associated with nodal strength only in P+CSO+ participants, within the subnetwork distinguishing them from controls. Together, these findings provide evidence for separable large-scale functional network signatures associated with pedohebephilic preference and offending history. Although the modest classification performance and absence of independent validation preclude diagnostic or forensic interpretation, the findings provide a neurobiological distinction for testing in larger, independent cohorts and connect pedohebephilia and pedohebephilic offending to broader research on self-referential processing, action selection, and impulse regulation.

## FUNDING STATEMENT

This research was funded through the Federal Ministry of Education and Research (BMBF): 01KR1205 to H.W., K. M. B., B. S., T. H. C. K., and M. W. as part of a multi-site research consortium on neural mechanisms underlying pedophilia and sexual offending against children (www.nemup.de) and by the grant from the German Research Foundation (DFG) to the ENIGMA task-based fMRI Working Group (DFG ER 724/4–1, WA 1539/11–1).

## DATA AVAILABILITY STATEMENT

Research data is not available as the participants did not consent to data sharing. Analysis code can be found at https://github.com/eminSerin/nemup-ml-paper.

## TECHNOLOGY USE DISCLOSURE

ChatGPT (OpenAI) and Claude (Anthropic) were used for proofreading (e.g., grammar, typos, integrity, and clarity checks) and as a coding assistant for the analysis scripts. All authors have read, corrected, and verified the information presented in this manuscript and its Supplementary Materials.

## CONFLICT OF INTERESTS

None.

## AUTHOR CONTRIBUTIONS

**Emin Serin:** Conceptualization, Methodology, Software, Validation, Formal analysis, Data Curation, Writing – Original Draft, Writing – Review & Editing, Visualization. **Kainat Raza Khan:** Methodology, Formal analysis, Data Curation, Writing – Original Draft, Writing – Review & Editing. **Tillman H. C. Krueger:** Writing – Review & Editing, Funding acquisition. **Christopher Sinke:** Writing – Review & Editing. **Klaus M. Beier:** Writing – Review & Editing, Funding acquisition. **Martin Walter**: Writing – Review & Editing, Funding acquisition. **Boris Schiffer:** Writing – Review & Editing, Funding acquisition. **Till Amelung:** Conceptualization, Writing – Original Draft, Writing – Review & Editing, Supervision. **Henrik Walter:** Resources, Writing – Review & Editing, Supervision, Funding acquisition. All authors ensure the accuracy of this work.

## FOOTNOTES

i According to DSM-5-TR and ICD-11, only the interest in prepubescent children can be diagnosed as pedophilic disorder if acted upon or accompanied by significant distress. The ICD-10 implicitly subsumed hebephilia under the criteria for pedophilia, since the age cutoff for preferred partners included early puberty. In contrast, current diagnostic systems such as DSM-5-TR and ICD-11 exclude this age from their criteria for pedophilia, making it classifiable only under “other specified paraphilic disorder”. Previous research on this matter therefore included pedohebephilic samples, which we also used.

## Notes

### Competing Interest Statement

The authors have declared no competing interest.

### Summary of Updates

The abstract was changed to free form; author affiliations, funding statements, and author contributions were updated.

## REFERENCES

1. Seto, M. C. Clinical and Conceptual Problems With Pedophilic Disorder in the DSM-5-TR. Arch. Sex. Behav. 51, 1833–1837 (2022).

2. Stephens, S., Seto, M. C., Cantor, J. M. & Lalumière, M. L. The Revised Screening Scale for Pedophilic Interests (SSPI-2) may be a measure of pedohebephilia. J. Sex. Med. 16, 1655–1663 (2019).

3. Bártová, K., Androvičová, R., Krejčová, L., Weiss, P. & Klapilová, K. The prevalence of paraphilic interests in the Czech population: Preference, arousal, the use of pornography, fantasy, and behavior. J. Sex Res. 58, 86–96 (2021).

4. Joyal, C. C. & Carpentier, J. The Prevalence of Paraphilic Interests and Behaviors in the General Population: A Provincial Survey. J. Sex Res. 54, 161–171 (2017).

5. Dombert, B. et al. How common is males’ self-reported sexual interest in prepubescent children? J. Sex Res. 53, 214–223 (2016).

6. Seto, M. C. Pedophilia and Sexual Offending against Children: Theory, Assessment, and Intervention. (American Psychological Association, Washington, DC, 2018). doi:10.1037/0000107-005.

7. Schiffer, B. et al. Structural brain abnormalities in the frontostriatal system and cerebellum in pedophilia. J. Psychiatr. Res. 41, 753–762 (2007).

8. Schiltz, K. et al. Brain pathology in pedophilic offenders: evidence of volume reduction in the right amygdala and related diencephalic structures. Arch. Gen. Psychiatry 64, 737 (2007).

9. Poeppl, T. B. et al. Association between brain structure and phenotypic characteristics in pedophilia. J. Psychiatr. Res. 47, 678–685 (2013).

10. Abé, C. et al. Brain structure and clinical profile point to neurodevelopmental factors involved in pedophilic disorder. Acta Psychiatr. Scand. 143, 363–374 (2021).

11. Schiffer, B. et al. Gray matter anomalies in pedophiles with and without a history of child sexual offending. Transl. Psychiatry 7, e1129 (2017).

12. Lett, T. A. et al. Multimodal neuroimaging measures and intelligence influence pedophile child sexual offense behavior. Eur. Neuropsychopharmacol. J. Eur. Coll. Neuropsychopharmacol. 28, 818–827 (2018).

13. Cantor, J. M. et al. Cerebral white matter deficiencies in pedophilic men. J. Psychiatr. Res. 42, 167–183 (2008).

14. Cantor, J. M. & Blanchard, R. White Matter Volumes in Pedophiles, Hebephiles, and Teleiophiles. Arch. Sex. Behav. 41, 749–752 (2012).

15. Tenbergen, G. et al. The Neurobiology and Psychology of Pedophilia: Recent Advances and Challenges. Front. Hum. Neurosci. 9, 344 (2015).

16. Mohnke, S. et al. Brain alterations in paedophilia: A critical review. Prog. Neurobiol. 122, 1–23 (2014).

17. Gerwinn, H. et al. The (in)consistency of changes in brain macrostructure in male paedophiles: A combined T1-weighted and diffusion tensor imaging study. J. Psychiatr. Res. 68, 246–253 (2015).

18. Cohen, L. J. & Galynker, I. I. Clinical features of pedophilia and implications for treatment. J. Psychiatr. Pract. 8, 276–289 (2002).

19. Kruger, T. H. C. & Schiffer, B. Neurocognitive and Personality Factors in Homo- and Heterosexual Pedophiles and Controls. J. Sex. Med. 8, 1650–1659 (2011).

20. Shehzad, Z. et al. The resting brain: unconstrained yet reliable. Cereb. Cortex N. Y. N 1SS1 19, 2209–2229 (2009).

21. Park, K. Y. et al. Mapping language function with task-based vs. resting-state functional MRI. PLOS ONE 15, e0236423 (2020).

22. Lee, M. H., Smyser, C. D. & Shimony, J. S. Resting-state fMRI: a review of methods and clinical applications. AJNR Am. J. Neuroradiol. 34, 1866–1872 (2013).

23. Zhang, Z. et al. Resting-state network alterations in depression: a comprehensive meta-analysis of functional connectivity. Psychol. Med. 55, e63 (2025).

24. Yang, X., Zhang, N. & Schrader, P. A study of brain networks for autism spectrum disorder classification using resting-state functional connectivity. Mach. Learn. Appl. 8, 100290 (2022).

25. Kärgel, C. et al. Diminished functional connectivity on the road to child sexual abuse in pedophilia. J. Sex. Med. 12, 783–795 (2015).

26. Cantor, J. M. et al. Independent Component Analysis of Resting-State Functional Magnetic Resonance Imaging in Pedophiles. J. Sex. Med. 13, 1546–1554 (2016).

27. Andersson, L. et al. Resting state functional connectivity in paedophilic disorder and degarelix treatment. Acta Neuropsychiatr. 38, e35 (2026).

28. Bressler, S. L. & Menon, V. Large-scale brain networks in cognition: emerging methods and principles. Trends Cogn. Sci. 14, 277–290 (2010).

29. Bullmore, E. T. & Bassett, D. S. Brain Graphs: Graphical Models of the Human Brain Connectome. Annu. Rev. Clin. Psychol. 7, 113–140 (2011).

30. Menon, V. Large-scale brain networks and psychopathology: a unifying triple network model. Trends Cogn. Sci. 15, 483–506 (2011).

31. Bassett, D. S. & Bullmore, E. T. Human brain networks in health and disease. Curr. Opin. Neurol. 22, 340–347 (2009).

32. Woo, C.-W., Chang, L. J., Lindquist, M. A. & Wager, T. D. Building better biomarkers: brain models in translational neuroimaging. Nat. Neurosci. 20, 365–377 (2017).

33. Friston, K. J. et al. Statistical parametric maps in functional imaging: A general linear approach. Hum. Brain Mapp. 2, 189–210 (1994).

34. Arbabshirani, M. R., Plis, S., Sui, J. & Calhoun, V. D. Single subject prediction of brain disorders in neuroimaging: Promises and pitfalls. NeuroImage 145, 137–165 (2017).

35. Ponseti, J. et al. Assessing paedophilia based on the haemodynamic brain response to face images. World J. Biol. Psychiatry 17, 39–46 (2016).

36. Ponseti, J. Assessment of Pedophilia Using Hemodynamic Brain Response to Sexual Stimuli. Arch. Gen. Psychiatry 69, 187 (2012).

37. Popovic, D. et al. Patterns of risk-Using machine learning and structural neuroimaging to identify pedophilic offenders. Front. Psychiatry 14, 1001085 (2023).

38. Serin, E., Zalesky, A., Matory, A., Walter, H. & Kruschwitz, J. D. NBS-Predict: A prediction-based extension of the network-based statistic. NeuroImage 244, 118625 (2021).

39. Turner, D. & Rettenberger, M. Neuropsychological functioning in child sexual abusers: A systematic review. Aggress. Violent Behav. 54, 101405 (2020).

40. Weidacker, K. et al. Interference inhibition in offending and non-offending pedophiles: A preliminary event-related fMRI study. Neuropsychologia 173, 108301 (2022).

41. Kärgel, C. et al. Evidence for superior neurobiological and behavioral inhibitory control abilities in non-offending as compared to offending pedophiles: Response Inhibition in Pedophilia. Hum. Brain Mapp. 38, 1092–1104 (2017).

42. World Health Organization. The ICD-10 Classification of Mental and Behavioural Disorders: Clinical Descriptions and Diagnostic Guidelines. vol. 1 (1992).

43. Stephens, S., Seto, M. C., Goodwill, A. M. & Cantor, J. M. Evidence of Construct Validity in the Assessment of Hebephilia. Arch. Sex. Behav. 46, 301–309 (2017).

44. Wechsler, D. Wechsler Adult Intelligence Scale--Fourth Edition. 10.1037/t15169-000 (2012).

45. First, M. B. Structured Clinical Interview for DSM-IV Axis I Disorders: SCID-I. (American Psychiatric Press; Biometrics Research Dept., New York State Psychiatric Institutute, Washington, DC, New York, N.Y., 1997).

46. User’s Guide for the Structured Clinical Interview for DSM-IV Axis II Personality Disorders: SCID-II. (American Psychiatric Publ, Arlington, Va, 1997).

47. Hamilton, M. THE ASSESSMENT OF ANXIETY STATES BY RATING. Br. J. Med. Psychol. 32, 50–55 (1959).

48. Hamilton, M. A rating scale for depression. J. Neurol. Neurosurg. Psychiatry 23, 56–62 (1960).

49. Kneer, J. et al. Diminished fronto-limbic functional connectivity in child sexual offenders. J. Psychiatr. Res. 108, 48–56 (2019).

50. Waller, L. et al. ENIGMA HALFpipe: interactive, reproducible, and efficient analysis for resting-state and task-based fMRI data. Hum. Brain Mapp. 43, 2727–2742 (2022).

51. Avants, B., Tustison, N. J. & Song, G. Advanced Normalization Tools: V1.0. Insight J. https://doi.org/10.54294/uvnhin (2009) doi:10.54294/uvnhin.

52. Pruim, R. H. R. et al. ICA-AROMA: A robust ICA-based strategy for removing motion artifacts from fMRI data. NeuroImage 112, 267–277 (2015).

53. Lindquist, M. A., Geuter, S., Wager, T. D. & Caffo, B. S. Modular preprocessing pipelines can reintroduce artifacts into fMRI data. Hum. Brain Mapp. 40, 2358–2376 (2019).

54. Power, J. D., Barnes, K. A., Snyder, A. Z., Schlaggar, B. L. & Petersen, S. E. Spurious but systematic correlations in functional connectivity MRI networks arise from subject motion. NeuroImage 59, 2142–2154 (2012).

55. Rolls, E. T., Huang, C.-C., Lin, C.-P., Feng, J. & Joliot, M. Automated anatomical labelling atlas 3. NeuroImage 206, 116189 (2020).

56. Zalesky, A., Fornito, A. & Bullmore, E. T. Network-based statistic: Identifying differences in brain networks. Neuroimage 53, 1197–1207 (2010).

57. Meskaldji, D. E. et al. Comparing connectomes across subjects and populations at different scales. Neuroimage https://doi.org/10.1016/j.neuroimage.2013.04.084 (2013) doi:10.1016/j.neuroimage.2013.04.084.

58. Snoek, L., Miletić, S. & Scholte, H. S. How to control for confounds in decoding analyses of neuroimaging data. NeuroImage 184, 741–760 (2019).

59. Ojala, M. & Garriga, G. C. Permutation tests for studying classifier performance. J Mach Learn Res 11, 1833–1863 (2010).

60. Benjamini, Y. & Hochberg, Y. Controlling the False Discovery Rate: A Practical and Powerful Approach to Multiple Testing. J. R. Stat. Soc. Ser. B Methodol. 57, 289–300 (1995).

61. Serin, E., Vaidya, N., Walter, H. & Kruschwitz, J. D. NBS-Predict: An Easy-to-Use Toolbox for Connectome-Based Machine Learning. in Methods for Analyzing Large Neuroimaging Datasets (eds Whelan, R. & Lemaître, H.) vol. 218 289–328 (Springer US, New York, NY, 2025).

62. Yeo, B. T. T. et al. The organization of the human cerebral cortex estimated by intrinsic functional connectivity. J. Neurophysiol. 106, 1125–1165 (2011).

63. Bryce, N. V. et al. Brain parcellation selection: An overlooked decision point with meaningful effects on individual differences in resting-state functional connectivity. NeuroImage 243, 118487 (2021).

64. Fan, L. et al. The Human Brainnetome Atlas: A New Brain Atlas Based on Connectional Architecture. Cereb. Cortex 26, 3508–3526 (2016).

65. Patton, J. H., Stanford, M. S. & Barratt, E. S. Factor structure of the Barratt impulsiveness scale. J. Clin. Psychol. 51, 768–774 (1995).

66. Penner, J. et al. Temporoparietal Junction Functional Connectivity in Early Schizophrenia and Major Depressive Disorder. Chronic Stress 2, 2470547018815232 (2018).

67. Yang, X. et al. Anhedonia correlates with abnormal functional connectivity of the superior temporal gyrus and the caudate nucleus in patients with first-episode drug-naive major depressive disorder. J. Affect. Disord. 218, 284–290 (2017).

68. Hu, L. et al. Disruption of resting-state functional connectivity of right posterior insula in adolescents and young adults with major depressive disorder. J. Affect. Disord. 257, 23–30 (2019).

69. Chen, C. et al. Decreased Cortical Folding of the Fusiform Gyrus and Its Hypoconnectivity with Sensorimotor Areas in Major Depressive Disorder. J. Affect. Disord. 295, 657–664 (2021).

70. Chen, C. P. et al. Diagnostic classification of intrinsic functional connectivity highlights somatosensory, default mode, and visual regions in autism. NeuroImage Clin. 8, 238–245 (2015).

71. Margari, A. et al. Autism spectrum disorder (ASD) and sexual offending: A systematic review. Neurosci Biobehav Rev 162, 105687 (2024).

72. Lätth, J., Joleby, M., McMahan, A., Luke, T. J. & Rahm, C. Child Sexual Abuse Material Users on the Darknet: Psychiatric Morbidities Related to Offence Behavior. Sex Abuse 10790632251347562, (2025).

73. Fraser, J. M., Babchishin, K. M. & Helmus, L. M. Emotional Congruence with Children: An Empirical Examination of Different Models in Men with a History of Sexually Offending Against Children. Sex. Abuse J. Res. Treat. 36, 546–571 (2024).

74. Finkelhor, D. & Araji, S. Explanations of pedophilia: A four factor model. J. Sex Res. 22, 145–161 (1986).

75. Cera, N. et al. The role of anterior and posterior insula in male genital response and in visual attention: an exploratory multimodal fMRI study. Sci. Rep. 10, 18463 (2020).

76. Kell, C. A., Kriegstein, K. von, Rösler, A., Kleinschmidt, A. & Laufs, H. The Sensory Cortical Representation of the Human Penis: Revisiting Somatotopy in the Male Homunculus. J. Neurosci. 25, 5984–5987 (2005).

77. Poeppl, T. B., Langguth, B., Laird, A. R. & Eickhoff, S. B. The functional neuroanatomy of male psychosexual and physiosexual arousal: A quantitative meta-analysis. Hum. Brain Mapp. 35, 1404–1421 (2014).

78. Panayiotopoulos, C. P. The Epilepsies: Seizures, Syndromes and Management: Based on the ILAE Classifications and Practice Parameter Guidelines. (Bladon Medical Publishing, Oxfordshire, UK, 2005).

79. Gattuso, J. J. et al. Default Mode Network Modulation by Psychedelics: A Systematic Review. Int. J. Neuropsychopharmacol. 26, 155–188 (2023).

80. McPhail, I. V. et al. Emotional Congruence with Children: Are Implicit and Explicit Child-Like Self-Concept and Attitude Toward Children Associated with Sexual Offending Against Children? Arch. Sex. Behav. 47, 2241–2254 (2018).

81. Hsu, K. J. & Bailey, J. M. Autopedophilia: Erotic-Target Identity Inversions in Men Sexually Attracted to Children. Psychol. Sci. 28, 115–123 (2017).

82. DeLong, M. R. Primate models of movement disorders of basal ganglia origin. Trends Neurosci. 13, 281–285 (1990).

83. Bublíková, I. et al. Structural Connectivity of the Substantia Nigra: A Comprehensive Review of Diffusion Imaging and Tractography Studies. Appl. Sci. 15, 7902 (2025).

84. Schnellbächer, G. J. et al. Aberrant brain activity in pedophilia links to receptor distribution, gene expression, and behavior. Nat. Ment. Health 1, 615–622 (2023).

85. Suchy, Y., Eastvold, A. D., Strassberg, D. S. & Franchow, E. I. Understanding processing speed weaknesses among pedophilic child molesters: Response style vs. Neuropathology. J. Abnorm. Psychol. 123, 273–285 (2014).

86. Massau, C. et al. Executive Functioning in Pedophilia and Child Sexual Offending. J. Int. Neuropsychol. Soc. 23, 460–470 (2017).

87. Eastvold, A., Suchy, Y. & Strassberg, D. Executive Function Profiles of Pedophilic and Nonpedophilic Child Molesters. J. Int. Neuropsychol. Soc. 17, 295–307 (2011).

88. Vabalas, A., Gowen, E., Poliakoff, E. & Casson, A. J. Machine learning algorithm validation with a limited sample size. PLOS ONE 14, e0224365 (2019).

89. Schnack, H. G. & Kahn, R. S. Detecting Neuroimaging Biomarkers for Psychiatric Disorders: Sample Size Matters. Front. Psychiatry 7, (2016).

90. Huang, W. & Shu, N. AI-powered integration of multimodal imaging in precision medicine for neuropsychiatric disorders. Cell Rep. Med. 6, 102132 (2025).

91. Du, L. et al. Unveiling the future: Advancements in MRI imaging for neurodegenerative disorders. Ageing Res. Rev. 95, 102230 (2024).

92. Marquand, A. F. et al. Conceptualizing mental disorders as deviations from normative functioning. Mol. Psychiatry 24, 1415–1424 (2019).

