## Supplementary Material for "Dissociating Resting-State Networks of Pedohebephilic Preference and Child Sexual Offending Using Network-Based Prediction"

† Equal supervision

**Supplementary Material**

### **Supplementary Methods**

**Nodal Strength**

Nodal strength is an analog of nodal degree in weighted networks, quantifying the sum of edge weights connected to a node. It identifies nodes (i.e., brain regions) with high connectivity, acting as network hubs (1). In an undirected network, nodal strength for node *i* is computed as:

$${NS}_{i}=\sum_{j\neq i} w_{ij}$$

where w_ij_ is the weight (i.e., relevancy score in NBS-Predict) of the edge linking nodes 𝑖 and 𝑗. In our study, high nodal strength indicates that the corresponding node plays a significant role as a network hub in the subnetwork identified as highly relevant for group classification.

**Barratt Impulsiveness Scale (BIS-11)**

The Barratt Impulsiveness Scale (BIS-11) is a widely used 30-item self-report questionnaire assessing three dimensions of impulsivity: attentional, motor, and non-planning impulsivity. Items are rated on a 4-point scale, yielding total scores ranging from 30 to 120, with higher scores indicating greater impulsivity. Psychometric properties of the English and German versions have been reported elsewhere (2,3). In the present study, the BIS-11 total score was used to index overall impulsivity.

### **Supplementary Figures**

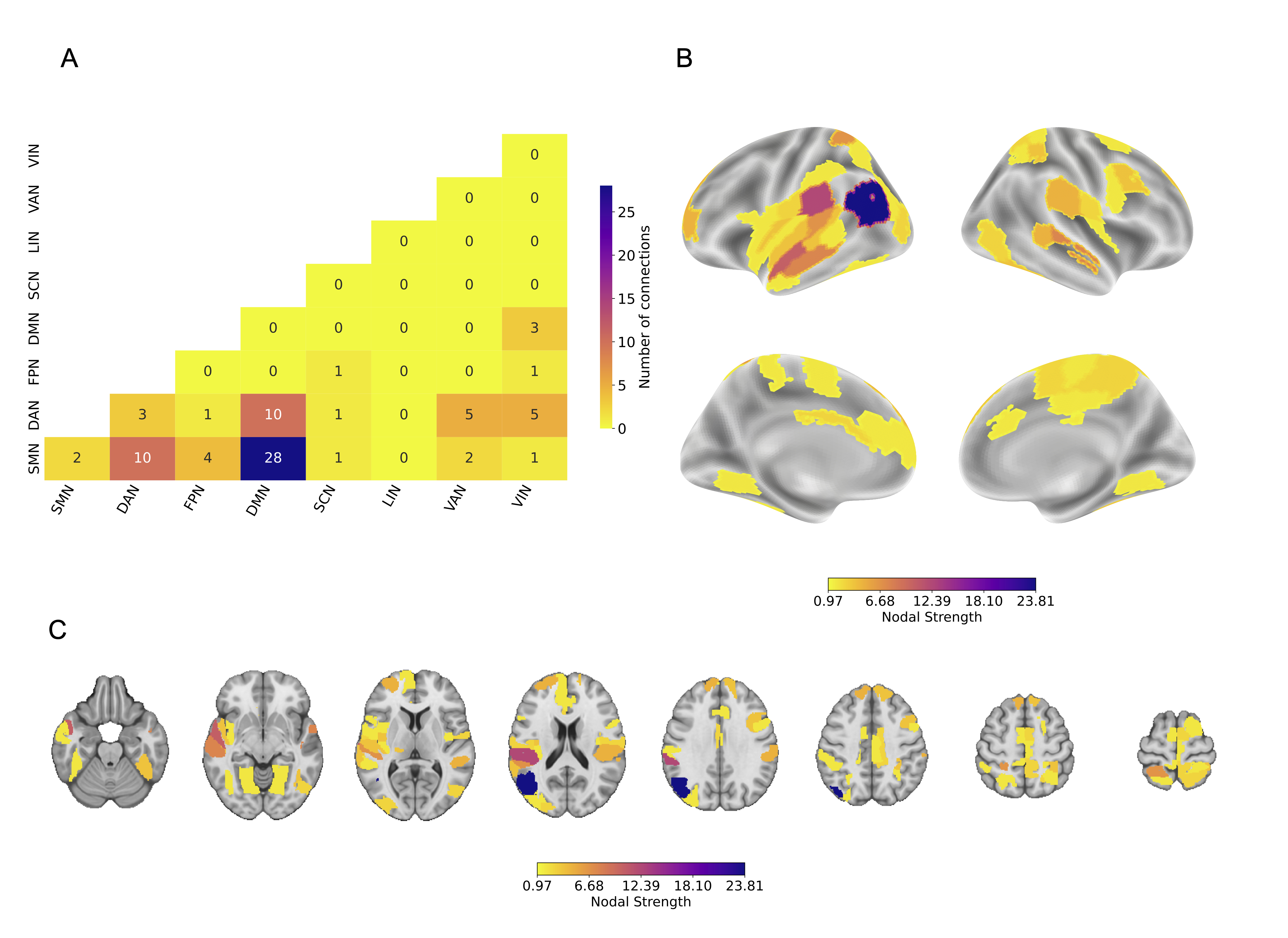

**Supplementary Figure 1.** Subnetwork of relevant decreased connections in P+CSO- (79 connections among 54 regions), classifying P+CSO- and P-CSO- groups using NBS-Predict. The feature set consists of participants' rsFC, computed as the pairwise correlation between the time series of brain regions delineated using the Brainnetome Atlas. A weight threshold of 0.97 was applied to visualize highly relevant connections for the classification. **A.** The heatmap illustrates the number of connections between canonical brain networks. The region-network mapping is given in Supplementary Table 1. **B.** and **C.** Nodal strength of brain regions in the subnetwork is displayed on the brain surface (fsaverage) and in volumetric space. The details of the nodal strength calculation are provided in the Supplementary Methods.

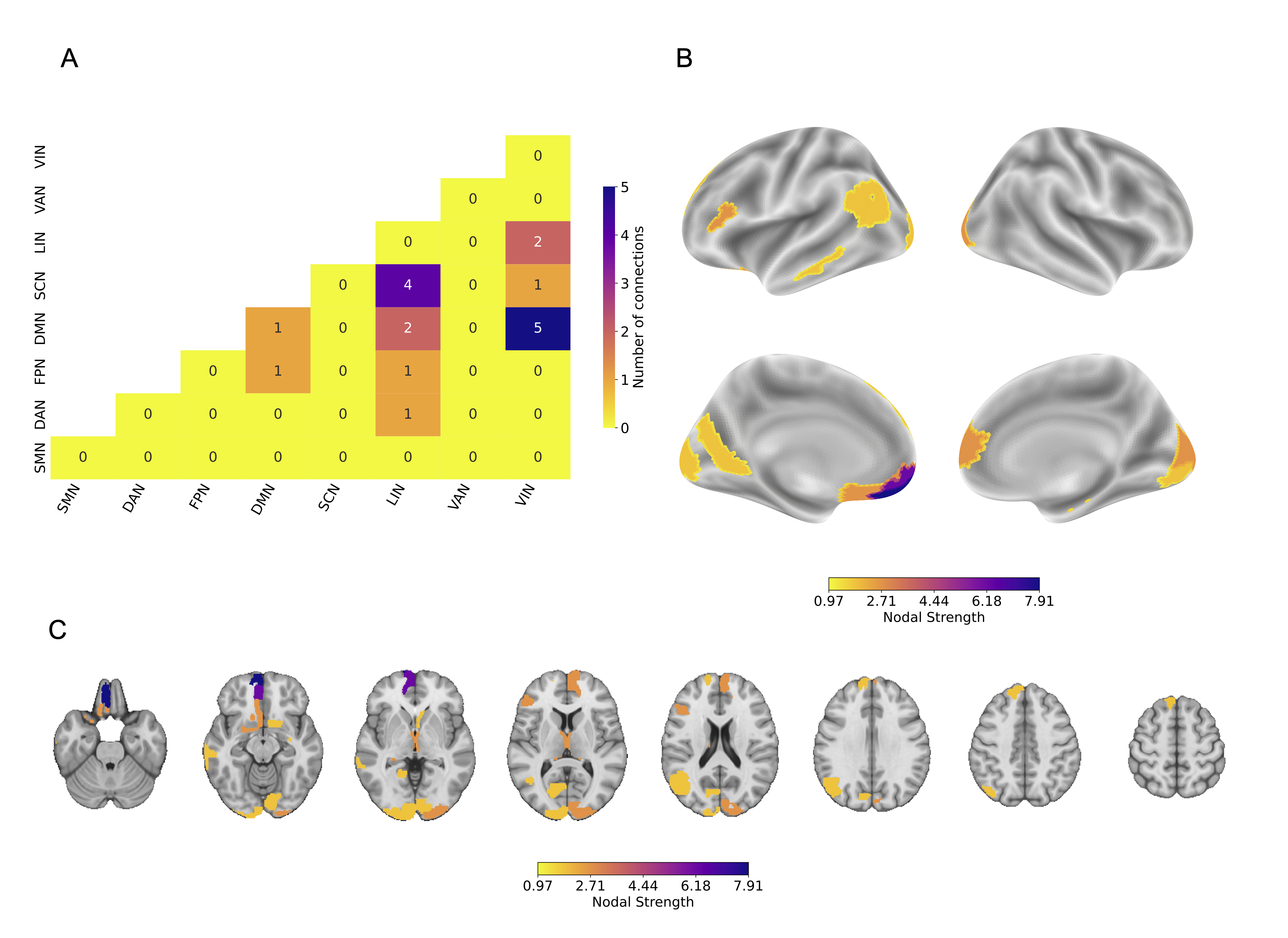

**Supplementary Figure 2.** Subnetwork of relevant decreased connections in P+CSO+ (18 connections among 17 regions), classifying P+CSO+ and P+CSO- groups using NBS-Predict. The feature set consists of participants' rsFC, computed as the pairwise correlation between the time series of brain regions delineated using the Brainnetome Atlas. A weight threshold of .97 was applied to visualize highly relevant connections for the classification. **A.** The heatmap illustrates the number of connections between canonical brain networks. The region-network mapping is given in Supplementary Table 1. **B.** and **C.** Nodal strength of brain regions in the subnetwork is displayed on the brain surface (fsaverage) and in volumetric space. The details of the nodal strength calculation are provided in the Supplementary Methods.

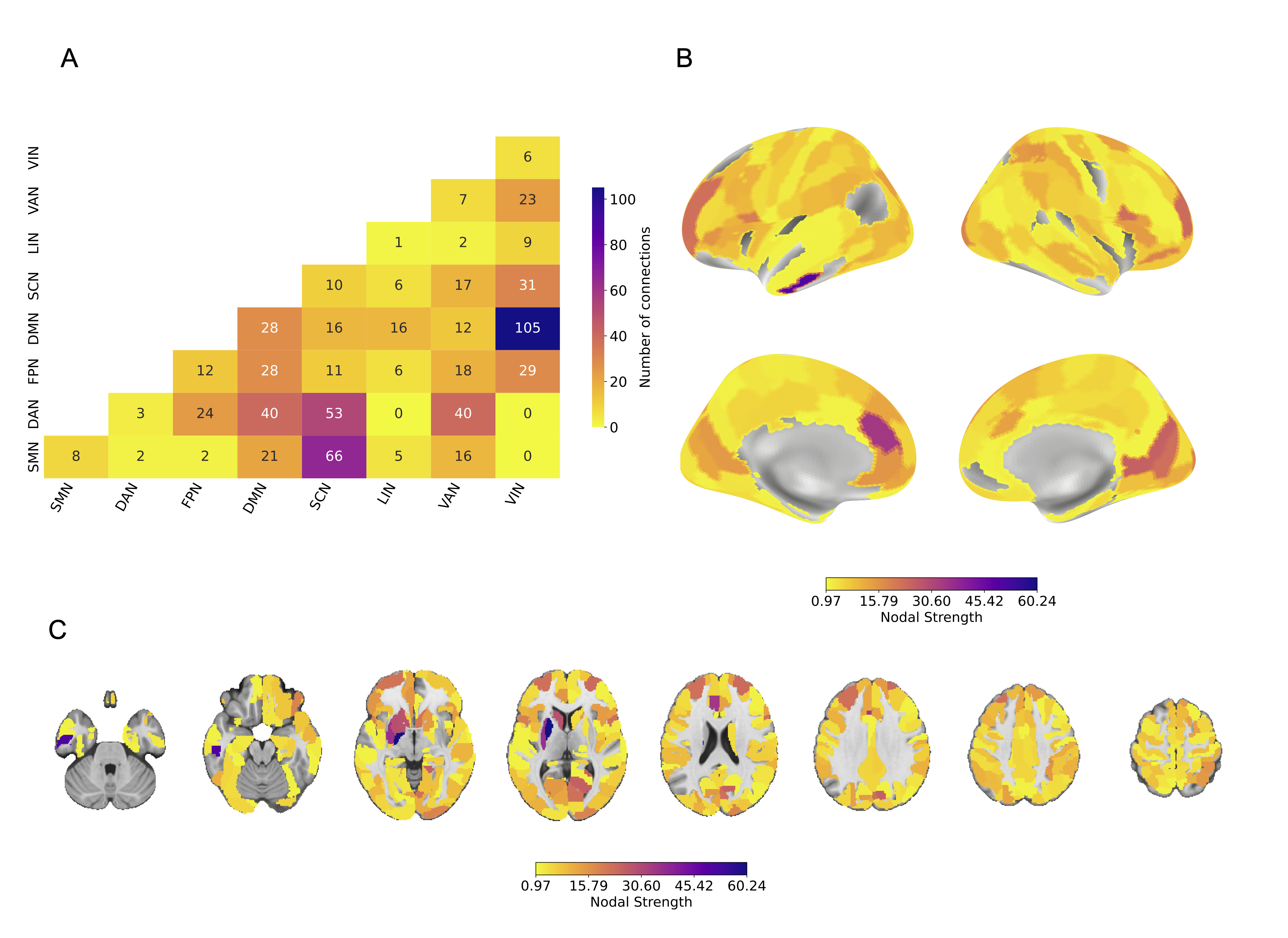

**Supplementary Figure 3.** Subnetwork of relevant decreased connections in P+CSO+ (673 connections among 190 regions), classifying P+CSO+ and P-CSO- groups using NBS-Predict. The feature set consists of participants' rsFC, computed as the pairwise correlation between the time series of brain regions delineated using the Brainnetome Atlas. A weight threshold of .97 was applied to visualize highly relevant connections for the classification. **A.** The heatmap illustrates the number of connections between canonical brain networks. The region-network mapping is given in Supplementary Table 1. **B.** and **C.** Nodal strength of brain regions in the subnetwork is displayed on the brain surface (fsaverage) and in volumetric space. The details of the nodal strength calculation are provided in the Supplementary Methods.

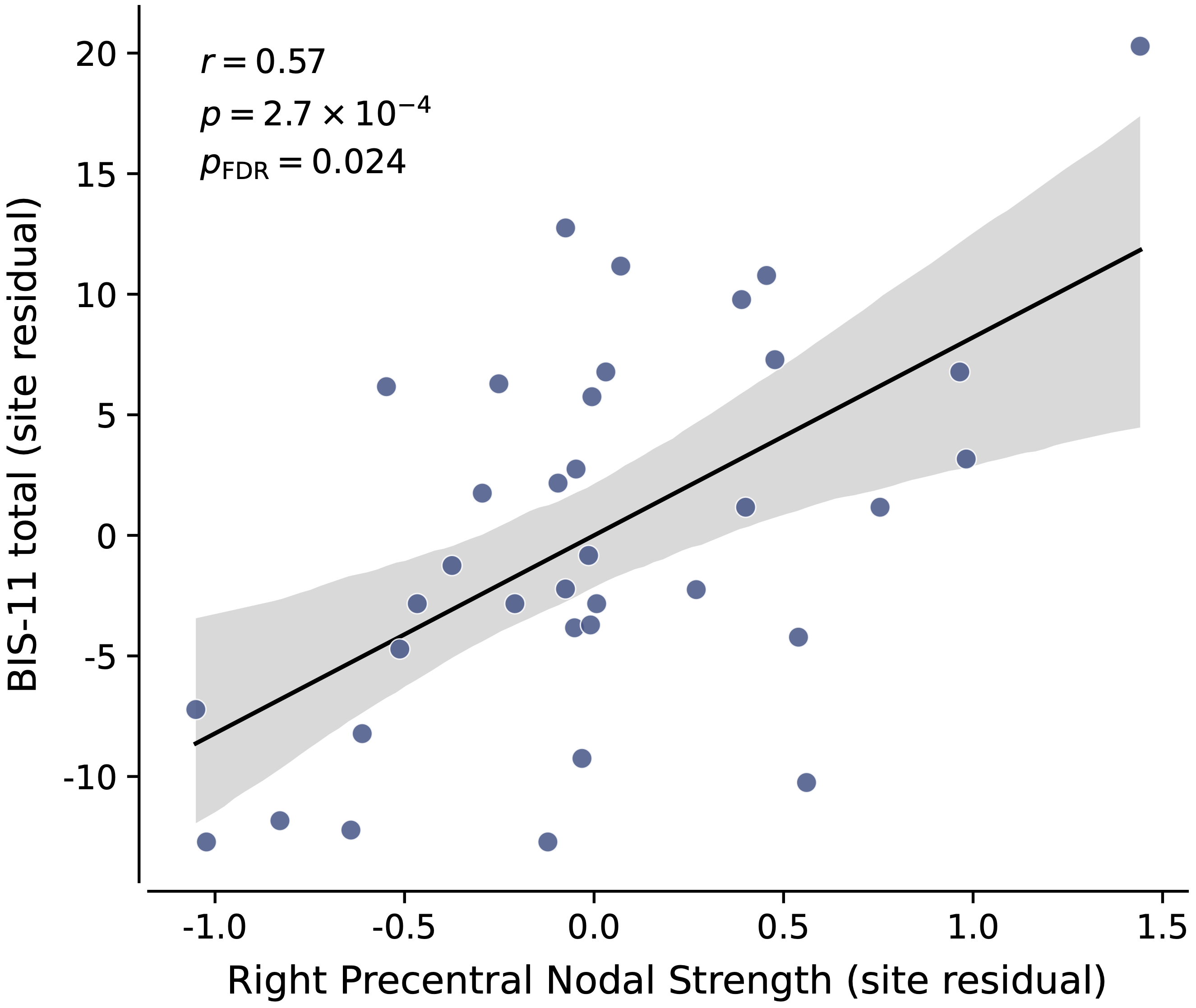

**Supplementary Figure 4.** Scatter plot of residualized BIS-11 total score versus nodal strength of the right precentral gyrus within the network identified by NBS-Predict (P-CSO- > P+CSO+). The site-corrected partial correlation value, as well as the uncorrected and FDR-corrected *p*-values, are annotated within the figure. The full results are given in Supplementary Table 11. No significant results were found for other contrasts (Supplementary Tables 5, 8).

### **Supplementary Tables**

| **Supplementary Table 1.** List of brain regions from AAL3 atlas (4) and the corresponding large-scale Yeo networks (5). | | | | | |
| --- | --- | --- | --- | --- | --- |
| **AAL Region** | **Network** | **AAL Region** | **Network** | **AAL Region** | **Network** |
| Precentral_L | SMN | Cuneus_R | VIS | Thal_VA_L | SCN |
| Precentral_R | SMN | Lingual_L | VIS | Thal_VA_R | SCN |
| Frontal_Sup_2_L | DMN | Lingual_R | VIS | Thal_VL_L | SCN |
| Frontal_Sup_2_R | DMN | Occipital_Sup_L | VIS | Thal_VL_R | SCN |
| Frontal_Mid_2_L | FPN | Occipital_Sup_R | VIS | Thal_VPL_L | SCN |
| Frontal_Mid_2_R | FPN | Occipital_Mid_L | VIS | Thal_VPL_R | SCN |
| Frontal_Inf_Oper_L | FPN | Occipital_Mid_R | VIS | Thal_IL_L | SCN |
| Frontal_Inf_Oper_R | FPN | Occipital_Inf_L | VIS | Thal_IL_R | SCN |
| Frontal_Inf_Tri_L | FPN | Occipital_Inf_R | VIS | Thal_Re_L | SCN |
| Frontal_Inf_Tri_R | FPN | Fusiform_L | VIS | Thal_Re_R | SCN |
| Frontal_Inf_Orb_2_L | DMN | Fusiform_R | VIS | Thal_MDm_L | SCN |
| Frontal_Inf_Orb_2_R | DMN | Postcentral_L | SMN | Thal_MDm_R | SCN |
| Rolandic_Oper_L | SMN | Postcentral_R | SMN | Thal_MDl_L | SCN |
| Rolandic_Oper_R | SMN | Parietal_Sup_L | DAN | Thal_MDl_R | SCN |
| Supp_Motor_Area_L | SMN | Parietal_Sup_R | DAN | Thal_LGN_L | SCN |
| Supp_Motor_Area_R | SMN | Parietal_Inf_L | DAN | Thal_LGN_R | SCN |
| Olfactory_L | LIM | Parietal_Inf_R | FPN | Thal_MGN_L | SCN |
| Olfactory_R | LIM | SupraMarginal_L | VAN | Thal_MGN_R | SCN |
| Frontal_Sup_Medial_L | DMN | SupraMarginal_R | VAN | Thal_PuI_L | SCN |
| Frontal_Sup_Medial_R | DMN | Angular_L | DMN | Thal_PuI_R | SCN |
| Frontal_Med_Orb_L | DMN | Angular_R | DMN | Thal_PuM_L | SCN |
| Frontal_Med_Orb_R | DMN | Precuneus_L | DMN | Thal_PuM_R | SCN |
| Rectus_L | LIM | Precuneus_R | DAN | Thal_PuA_L | SCN |
| Rectus_R | LIM | Paracentral_Lobule_L | SMN | Thal_PuA_R | SCN |
| OFCmed_L | LIM | Paracentral_Lobule_R | SMN | Thal_PuL_L | SCN |
| OFCmed_R | LIM | Caudate_L | SCN | Thal_PuL_R | SCN |
| OFCant_L | LIM | Caudate_R | SCN | ACC_sub_L | DMN |
| OFCant_R | FPN | Putamen_L | SCN | ACC_sub_R | DMN |
| OFCpost_L | LIM | Putamen_R | SCN | ACC_pre_L | DMN |
| OFCpost_R | LIM | Pallidum_L | SCN | ACC_pre_R | DMN |
| OFClat_L | DMN | Pallidum_R | SCN | ACC_sup_L | VAN |
| OFClat_R | DMN | Heschl_L | SMN | ACC_sup_R | VAN |
| Insula_L | VAN | Heschl_R | SMN | N_Acc_L | SCN |
| Insula_R | VAN | Temporal_Sup_L | SMN | N_Acc_R | SCN |
| Cingulate_Mid_L | VAN | Temporal_Sup_R | SMN | VTA_L | SCN |
| Cingulate_Mid_R | VAN | Temporal_Pole_Sup_L | LIM | VTA_R | SCN |
| Cingulate_Post_L | DMN | Temporal_Pole_Sup_R | LIM | SN_pc_L | SCN |
| Cingulate_Post_R | DMN | Temporal_Mid_L | DMN | SN_pc_R | SCN |
| Hippocampus_L | LIM | Temporal_Mid_R | DMN | SN_pr_L | SCN |
| Hippocampus_R | LIM | Temporal_Pole_Mid_L | LIM | SN_pr_R | SCN |
| ParaHippocampal_L | LIM | Temporal_Pole_Mid_R | LIM | Red_N_L | SCN |
| ParaHippocampal_R | LIM | Temporal_Inf_L | LIM | Red_N_R | SCN |
| Amygdala_L | LIM | Temporal_Inf_R | LIM | LC_L | SCN |
| Amygdala_R | LIM | Thal_AV_L | SCN | LC_R | SCN |
| Calcarine_L | VIS | Thal_AV_R | SCN | Raphe_D | SCN |
| Calcarine_R | VIS | Thal_LP_L | SCN | Raphe_M | SCN |
| Cuneus_L | VIS | Thal_LP_R | SCN |  |  |
| **Note:** The anatomical regions in the AAL3 atlas (4) were mapped onto 7 canonical resting-state networks (5): VIS, visual network; LIM, limbic network; SMN, somatomotor network; FPN, frontoparietal network; DAN, dorsal attention network; DMN, default mode network; and VAN, ventral attention (or salience) network. The remaining subcortical regions are assigned to the subcortical network (SCN). | | | | | |

| **Supplementary Table 2.** Results of all three pairwise classifications using NBS-Predict using Brainnetome Atlas. | | | | | |
| --- | --- | --- | --- | --- | --- |
|  | *Method* | *Balanced Accuracy* | *p_FDR_* | *AUC-ROC* | *F1* |
| **P-CSO- > P+CSO-** |  |  |  |  |  |
|  | **SVC** | **.589** | **.033** | .597 | .648 |
|  | **Logistic Regression** | **.558** | **.031** | .536 | .707 |
|  | **LDA** | .549 | .281 | .531 | .581 |
| **P+CSO- > P+CSO+** |  |  |  |  |  |
|  | **SVC** | **.640** | **.031** | .626 | .584 |
|  | **Logistic Regression** | **.613** | **.033** | .618 | .552 |
|  | **LDA** | **.638** | **.031** | .605 | .564 |
| **P-CSO- > P+CSO+** |  |  |  |  |  |
|  | **SVC** | **.650** | **.018** | .652 | .714 |
|  | **Logistic Regression** | **.627** | **.031** | .626 | .753 |
|  | **LDA** | .543 | .180 | .640 | .696 |
| **Note:** P+CSO+, individuals with pedohebephilic preference and a history of child sexual offending; P+CSO-, individuals with pedohebephilic preference without a history of child sexual offending; P-CSO-, non-pedohebephilic controls without a history of child sexual offending; SVC, support vector classifier; LDA, linear discriminant analysis; AUC-ROC, area under the receiver operating characteristic curve. A 50 repeated 7x7 nested cross-validation was used for model evaluation and hyperparameter optimization. Age, site, HAMD, and HAMA scores were controlled for. P-values were computed using permutation testing (1,000 permutations). FDR correction was applied across multiple classifications and classifiers (9 in total for subgroup classification). The significant classifications (*p* < .05) are highlighted. Classifications using reverse subnetworks (e.g., P-CSO- < P+CSO+) failed as no subnetworks survived feature selection, and, thus, the model could not be trained. | | | | | |

| **Supplementary Table 3.** Nodal degree and nodal strength of brain regions within the most relevant network identified by NBS-Predict (P-CSO- > P+CSO-; weight threshold ≥ .97). | | |
| --- | --- | --- |
| **AAL Label** | **Nodal Degree** | **Nodal Strength** |
| Temporal_Mid_L | 10 | 9.904 |
| SupraMarginal_L | 7 | 6.919 |
| Rolandic_Oper_L | 4 | 3.983 |
| Temporal_Mid_R | 4 | 3.969 |
| Angular_L | 4 | 3.959 |
| SupraMarginal_R | 3 | 2.948 |
| Frontal_Inf_Orb_2_L | 2 | 1.962 |
| Rolandic_Oper_R | 2 | 1.983 |
| Temporal_Sup_L | 2 | 1.990 |
| Paracentral_Lobule_L | 2 | 1.977 |
| Frontal_Sup_Medial_L | 1 | .987 |
| Frontal_Sup_Medial_R | 1 | .973 |
| Postcentral_R | 1 | .977 |
| Lingual_R | 1 | .987 |
| Occipital_Mid_R | 1 | .973 |
| Postcentral_L | 1 | .993 |
| Heschl_L | 1 | .996 |
| Paracentral_Lobule_R | 1 | .975 |
| Temporal_Pole_Sup_L | 1 | .976 |
| Temporal_Inf_L | 1 | .996 |
| **Note:** Nodal strength was computed from the edge weight of connections linking the corresponding brain regions. | | |

| **Supplementary Table 4.** Connections within robust connected network identified by NBS-Predict (P-CSO- > P+CSO-; weight threshold ≥ .97). | | |
| --- | --- | --- |
| **AAL Region A** | **AAL Region B** | **Weight** |
| Rolandic_Oper_L | Temporal_Mid_L | 1.000 |
| SupraMarginal_L | Angular_L | 1.000 |
| SupraMarginal_L | Temporal_Mid_L | 1.000 |
| SupraMarginal_L | Temporal_Mid_R | 1.000 |
| Temporal_Sup_L | Temporal_Mid_L | 1.000 |
| Rolandic_Oper_R | Temporal_Mid_L | .996 |
| Heschl_L | Temporal_Mid_L | .996 |
| Rolandic_Oper_L | Temporal_Inf_L | .996 |
| SupraMarginal_R | Temporal_Mid_L | .994 |
| Rolandic_Oper_L | Angular_L | .994 |
| Postcentral_L | Temporal_Mid_L | .993 |
| Rolandic_Oper_L | Temporal_Mid_R | .993 |
| Angular_L | Temporal_Sup_L | .990 |
| Paracentral_Lobule_L | Temporal_Mid_R | .989 |
| Lingual_R | Paracentral_Lobule_L | .987 |
| Frontal_Sup_Medial_L | SupraMarginal_L | .987 |
| Rolandic_Oper_R | Temporal_Mid_R | .986 |
| Frontal_Inf_Orb_2_L | SupraMarginal_L | .983 |
| Frontal_Inf_Orb_2_L | SupraMarginal_R | .979 |
| Postcentral_R | Temporal_Mid_L | .977 |
| SupraMarginal_L | Temporal_Pole_Sup_L | .976 |
| Paracentral_Lobule_R | Temporal_Mid_L | .975 |
| SupraMarginal_R | Angular_L | .975 |
| Frontal_Sup_Medial_R | SupraMarginal_L | .973 |
| Occipital_Mid_R | Temporal_Mid_L | .973 |

| **Supplementary Table 5.** Partial correlations between BIS-11 total scores and nodal strength of regions within the relevant network, identified by NBS-Prediction, in P+ (P-CSO- > P+CSO-; weight threshold ≥ .97). | | | |
| --- | --- | --- | --- |
| **AAL Region** | ***r*** | ***p*** | ***p_FDR_*** |
| Temporal_Mid_L | .247 | .166 | .371 |
| SupraMarginal_L | .253 | .156 | .371 |
| Rolandic_Oper_L | .081 | .656 | .729 |
| Temporal_Mid_R | .162 | .369 | .553 |
| Angular_L | .088 | .624 | .729 |
| SupraMarginal_R | .280 | .115 | .371 |
| Frontal_Inf_Orb_2_L | .236 | .186 | .371 |
| Rolandic_Oper_R | .123 | .494 | .618 |
| Temporal_Sup_L | .066 | .715 | .752 |
| Paracentral_Lobule_L | .283 | .110 | .371 |
| Frontal_Sup_Medial_L | .221 | .216 | .394 |
| Frontal_Sup_Medial_R | .245 | .170 | .371 |
| Postcentral_R | .250 | .160 | .371 |
| Lingual_R | .329 | .062 | .371 |
| Occipital_Mid_R | .379 | .030 | .371 |
| Postcentral_L | .156 | .387 | .553 |
| Heschl_L | .138 | .443 | .590 |
| Paracentral_Lobule_R | .252 | .157 | .371 |
| Temporal_Pole_Sup_L | .211 | .239 | .399 |
| Temporal_Inf_L | .056 | .758 | .758 |
| **Note:** Dummy-coded site variables were corrected in partial correlation. FDR correction was applied across brain regions within each contrast. Significant correlations are highlighted. | | | |

| **Supplementary Table 6.** Nodal degree and nodal strength of brain regions within the most relevant network identified by NBS-Predict (P+CSO- > P+CSO+; weight threshold ≥ .97). | | |
| --- | --- | --- |
| **AAL Label** | **Nodal Degree** | **Nodal Strength** |
| SN_pr_L | 6 | 5.898 |
| ACC_pre_L | 2 | 1.965 |
| Calcarine_L | 2 | 1.969 |
| OFCmed_L | 1 | .975 |
| Calcarine_R | 1 | .982 |
| Occipital_Mid_R | 1 | .990 |
| Occipital_Mid_L | 1 | .997 |
| Parietal_Sup_L | 1 | .970 |
| Occipital_Inf_R | 1 | .979 |
| **Note:** Nodal strength was computed from the edge weight of connections linking the corresponding brain regions. | | |

| **Supplementary Table 7.** Connections within robust connected network identified by NBS-Predict (P+CSO- > P+CSO+; weight threshold ≥ .97). | | |
| --- | --- | --- |
| **AAL Region A** | **AAL Region B** | **Weight** |
| OFCmed_L | SN_pr_L | .975 |
| Calcarine_L | ACC_pre_L | .982 |
| Calcarine_L | SN_pr_L | .987 |
| Calcarine_R | ACC_pre_L | .982 |
| Occipital_Mid_L | SN_pr_L | .997 |
| Occipital_Mid_R | SN_pr_L | .990 |
| Occipital_Inf_R | SN_pr_L | .979 |
| Parietal_Sup_L | SN_pr_L | .970 |

| **Supplementary Table 8.** Partial correlations between BIS-11 total scores and nodal strength of regions within the relevant network, identified by NBS-Prediction, in P+ (P+CSO- > P+CSO+; weight threshold ≥ .97). | | | |
| --- | --- | --- | --- |
| **AAL Region** | ***r*** | ***p*** | ***p_FDR_*** |
| SN_pr_L | -.341 | .052 | .122 |
| ACC_pre_L | .136 | .450 | .450 |
| Calcarine_L | -.198 | .270 | .347 |
| OFCmed_L | -.148 | .411 | .450 |
| Calcarine_R | .209 | .244 | .347 |
| Occipital_Mid_R | -.352 | .044 | .122 |
| Occipital_Mid_L | -.338 | .054 | .122 |
| Parietal_Sup_L | -.345 | .049 | .122 |
| Occipital_Inf_R | -.257 | .149 | .268 |
| **Note:** Dummy-coded site variables were corrected in partial correlation. FDR correction was applied across brain regions within each contrast. Significant correlations are highlighted. | | | |

| **Supplementary Table 9.** Nodal degree and nodal strength of brain regions within the most relevant network identified by NBS-Predict (P-CSO- > P+CSO+; weight threshold ≥ .97). | | |
| --- | --- | --- |
| **AAL Label** | **Nodal Degree** | **Nodal Strength** |
| Pallidum_L | 31 | 30.634 |
| Putamen_L | 21 | 20.693 |
| Calcarine_R | 14 | 13.903 |
| ACC_pre_L | 12 | 11.951 |
| ACC_sup_L | 10 | 9.904 |
| OFCant_R | 10 | 9.861 |
| Lingual_R | 10 | 9.901 |
| Frontal_Mid_2_L | 8 | 7.906 |
| Precentral_L | 8 | 7.924 |
| SN_pr_R | 8 | 7.900 |
| Red_N_R | 7 | 6.881 |
| N_Acc_R | 6 | 5.896 |
| Occipital_Mid_L | 6 | 5.971 |
| SupraMarginal_L | 5 | 4.915 |
| Hippocampus_R | 5 | 4.938 |
| Occipital_Inf_R | 5 | 4.938 |
| Fusiform_L | 5 | 4.930 |
| Calcarine_L | 5 | 4.940 |
| Frontal_Med_Orb_L | 5 | 4.973 |
| Frontal_Sup_Medial_L | 5 | 4.956 |
| SupraMarginal_R | 5 | 4.905 |
| Temporal_Mid_R | 5 | 4.937 |
| Frontal_Sup_Medial_R | 4 | 3.944 |
| Frontal_Sup_2_L | 4 | 3.960 |
| Precentral_R | 4 | 3.956 |
| ACC_pre_R | 4 | 3.969 |
| ACC_sub_R | 4 | 3.925 |
| Parietal_Sup_R | 4 | 3.920 |
| OFClat_R | 4 | 3.935 |
| Frontal_Sup_2_R | 3 | 2.981 |
| Frontal_Inf_Oper_R | 3 | 2.954 |
| ACC_sub_L | 3 | 2.987 |
| Temporal_Pole_Sup_R | 3 | 2.969 |
| SN_pc_R | 3 | 2.957 |
| ACC_sup_R | 3 | 2.976 |
| Angular_R | 3 | 2.963 |
| Supp_Motor_Area_L | 3 | 2.961 |
| Frontal_Inf_Orb_2_R | 3 | 2.957 |
| Supp_Motor_Area_R | 3 | 2.966 |
| Olfactory_R | 3 | 2.947 |
| Frontal_Med_Orb_R | 3 | 2.962 |
| Parietal_Sup_L | 3 | 2.975 |
| ParaHippocampal_R | 3 | 2.955 |
| Cingulate_Mid_L | 3 | 2.958 |
| Lingual_L | 3 | 2.956 |
| Postcentral_R | 3 | 2.968 |
| Caudate_L | 2 | 1.975 |
| Thal_PuL_R | 2 | 1.983 |
| Angular_L | 2 | 1.988 |
| Frontal_Inf_Oper_L | 2 | 1.969 |
| Frontal_Mid_2_R | 2 | 1.993 |
| Rolandic_Oper_R | 2 | 1.990 |
| N_Acc_L | 2 | 1.945 |
| Raphe_D | 2 | 1.977 |
| Temporal_Sup_R | 2 | 1.966 |
| Fusiform_R | 2 | 1.963 |
| Frontal_Inf_Tri_R | 2 | 1.979 |
| Postcentral_L | 2 | 1.978 |
| Occipital_Mid_R | 2 | 1.993 |
| Cingulate_Post_L | 2 | 1.975 |
| Parietal_Inf_R | 2 | 1.972 |
| Insula_L | 2 | 1.975 |
| Rolandic_Oper_L | 1 | .979 |
| Frontal_Inf_Tri_L | 1 | .987 |
| Frontal_Inf_Orb_2_L | 1 | .990 |
| OFCant_L | 1 | 1.000 |
| OFCmed_R | 1 | 1.000 |
| OFCmed_L | 1 | .979 |
| Rectus_R | 1 | .984 |
| OFCpost_R | 1 | .983 |
| Insula_R | 1 | .971 |
| Hippocampus_L | 1 | .993 |
| Amygdala_R | 1 | .972 |
| Parietal_Inf_L | 1 | .985 |
| Occipital_Inf_L | 1 | 1.000 |
| Occipital_Sup_L | 1 | .980 |
| OFClat_L | 1 | .997 |
| Cingulate_Mid_R | 1 | .972 |
| Cingulate_Post_R | 1 | .974 |
| Paracentral_Lobule_R | 1 | .980 |
| Precuneus_R | 1 | .990 |
| Temporal_Mid_L | 1 | .985 |
| Temporal_Pole_Sup_L | 1 | .971 |
| Putamen_R | 1 | .989 |
| Caudate_R | 1 | .986 |
| Precuneus_L | 1 | .997 |
| Temporal_Inf_R | 1 | .979 |
| Thal_PuM_L | 1 | .987 |
| Raphe_M | 1 | .997 |
| **Note:** Nodal strength was computed from the edge weight of connections linking the corresponding brain regions. | | |

| **Supplementary Table 10.** Connections within robust connected network identified by NBS-Predict (P-CSO- > P+CSO+; weight threshold ≥ .97). | | |
| --- | --- | --- |
| **AAL Region A** | **AAL Region B** | **Weight** |
| Precentral_L | Frontal_Mid_2_L | 1.000 |
| Precentral_L | Putamen_L | 1.000 |
| Precentral_L | Pallidum_L | 1.000 |
| Frontal_Sup_2_L | Pallidum_L | 1.000 |
| Frontal_Sup_2_R | Pallidum_L | 1.000 |
| Frontal_Mid_2_L | Insula_L | 1.000 |
| Frontal_Inf_Orb_2_R | Temporal_Mid_R | 1.000 |
| Rolandic_Oper_R | Pallidum_L | 1.000 |
| Frontal_Sup_Medial_L | Calcarine_R | 1.000 |
| Frontal_Med_Orb_L | Calcarine_R | 1.000 |
| Frontal_Med_Orb_L | Lingual_R | 1.000 |
| OFCmed_R | Red_N_R | 1.000 |
| OFCant_L | SN_pr_R | 1.000 |
| Hippocampus_R | ACC_sub_R | 1.000 |
| Calcarine_R | ACC_sup_L | 1.000 |
| Lingual_L | ACC_pre_L | 1.000 |
| Lingual_R | ACC_pre_L | 1.000 |
| Occipital_Mid_L | ACC_pre_L | 1.000 |
| Occipital_Inf_L | ACC_pre_L | 1.000 |
| Occipital_Inf_R | ACC_sup_L | 1.000 |
| Fusiform_L | ACC_pre_L | 1.000 |
| Red_N_R | Raphe_D | 1.000 |
| Calcarine_R | Angular_R | .997 |
| Parietal_Sup_L | SN_pc_R | .997 |
| Supp_Motor_Area_R | Putamen_L | .997 |
| OFClat_L | Calcarine_R | .997 |
| Occipital_Mid_L | ACC_pre_R | .997 |
| Occipital_Mid_R | ACC_pre_L | .997 |
| Lingual_R | ACC_pre_R | .997 |
| Frontal_Sup_2_L | Calcarine_R | .997 |
| Frontal_Sup_2_R | OFCant_R | .997 |
| Postcentral_R | Putamen_L | .997 |
| ACC_sub_L | Raphe_M | .997 |
| Frontal_Mid_2_L | Pallidum_L | .997 |
| Frontal_Mid_2_R | Pallidum_L | .997 |
| Frontal_Sup_Medial_L | Pallidum_L | .997 |
| Cingulate_Mid_L | Pallidum_L | .997 |
| Hippocampus_R | N_Acc_R | .997 |
| Occipital_Inf_R | ACC_sup_R | .997 |
| Parietal_Inf_R | Pallidum_L | .997 |
| SupraMarginal_R | Pallidum_L | .997 |
| Precuneus_L | Pallidum_L | .997 |
| Pallidum_L | Temporal_Pole_Sup_R | .997 |
| Frontal_Mid_2_R | OFCant_R | .996 |
| Frontal_Inf_Tri_R | OFCant_R | .996 |
| Supp_Motor_Area_L | SupraMarginal_L | .996 |
| Calcarine_R | ACC_sup_R | .996 |
| Occipital_Mid_L | ACC_sup_L | .996 |
| Occipital_Mid_R | ACC_sup_L | .996 |
| ACC_sub_L | SN_pr_R | .996 |
| Calcarine_R | ACC_pre_L | .996 |
| Frontal_Sup_Medial_R | Lingual_R | .996 |
| Frontal_Med_Orb_L | Calcarine_L | .996 |
| Frontal_Med_Orb_L | Occipital_Mid_L | .996 |
| Calcarine_R | Angular_L | .996 |
| Thal_PuL_R | ACC_sub_L | .994 |
| OFCant_R | SN_pr_R | .994 |
| Precentral_R | Putamen_L | .994 |
| Postcentral_L | Putamen_L | .994 |
| Frontal_Mid_2_L | Putamen_L | .993 |
| Calcarine_L | ACC_pre_L | .993 |
| Occipital_Inf_R | ACC_pre_L | .993 |
| Fusiform_L | ACC_sup_L | .993 |
| Precentral_L | Caudate_L | .993 |
| Hippocampus_L | ACC_pre_L | .993 |
| Lingual_R | ACC_sup_L | .993 |
| Precentral_L | ACC_pre_L | .993 |
| Angular_L | Pallidum_L | .993 |
| Frontal_Sup_Medial_L | Occipital_Mid_L | .993 |
| Frontal_Sup_Medial_L | Lingual_R | .992 |
| Frontal_Med_Orb_R | Lingual_R | .992 |
| Postcentral_R | Pallidum_L | .992 |
| ParaHippocampal_R | Putamen_L | .991 |
| Calcarine_L | ACC_sup_L | .990 |
| Calcarine_R | ACC_pre_R | .990 |
| Parietal_Sup_L | Putamen_L | .990 |
| Frontal_Sup_2_L | Occipital_Mid_L | .990 |
| Precuneus_R | Pallidum_L | .990 |
| Frontal_Sup_Medial_R | Calcarine_R | .990 |
| Rolandic_Oper_R | Putamen_L | .990 |
| Frontal_Inf_Orb_2_L | OFClat_R | .990 |
| Frontal_Mid_2_L | OFCant_R | .989 |
| Thal_PuL_R | SN_pr_R | .989 |
| Supp_Motor_Area_R | Putamen_R | .989 |
| Cingulate_Post_L | Calcarine_R | .989 |
| ParaHippocampal_R | N_Acc_R | .989 |
| Frontal_Inf_Oper_R | SupraMarginal_L | .988 |
| Precentral_R | Pallidum_L | .988 |
| Parietal_Sup_L | Pallidum_L | .988 |
| Precentral_R | OFClat_R | .988 |
| Frontal_Inf_Oper_L | Frontal_Inf_Tri_L | .987 |
| Olfactory_R | Thal_PuM_L | .987 |
| Parietal_Sup_R | Putamen_L | .987 |
| Putamen_L | Temporal_Pole_Sup_R | .987 |
| Frontal_Med_Orb_R | SN_pr_R | .987 |
| Cingulate_Mid_L | Lingual_R | .986 |
| Hippocampus_R | ACC_pre_L | .986 |
| Temporal_Sup_R | Temporal_Mid_R | .986 |
| Olfactory_R | Cingulate_Post_L | .986 |
| Pallidum_L | Temporal_Mid_R | .986 |
| Precentral_L | Caudate_R | .986 |
| Precentral_R | N_Acc_R | .986 |
| Frontal_Inf_Orb_2_R | Temporal_Pole_Sup_R | .986 |
| Supp_Motor_Area_L | Pallidum_L | .986 |
| Calcarine_L | ACC_pre_R | .985 |
| Pallidum_L | Temporal_Mid_L | .985 |
| Angular_R | Temporal_Mid_R | .985 |
| Postcentral_L | Pallidum_L | .985 |
| Parietal_Inf_L | Pallidum_L | .985 |
| Frontal_Sup_2_R | Calcarine_R | .984 |
| SupraMarginal_L | Putamen_L | .984 |
| Rectus_R | SN_pr_R | .984 |
| Frontal_Sup_Medial_R | OFCant_R | .984 |
| Fusiform_R | Putamen_L | .983 |
| Frontal_Med_Orb_R | Hippocampus_R | .983 |
| Fusiform_L | ACC_sup_R | .983 |
| Frontal_Inf_Oper_R | SupraMarginal_R | .983 |
| Frontal_Inf_Oper_R | OFCant_R | .983 |
| OFCpost_R | Red_N_R | .983 |
| Frontal_Inf_Tri_R | OFClat_R | .983 |
| Fusiform_L | Caudate_L | .982 |
| Frontal_Med_Orb_L | Lingual_L | .982 |
| Frontal_Mid_2_L | Frontal_Inf_Oper_L | .982 |
| Angular_R | Pallidum_L | .981 |
| Precentral_L | ACC_sub_R | .980 |
| SupraMarginal_R | Temporal_Mid_R | .980 |
| Parietal_Sup_R | SN_pc_R | .980 |
| Supp_Motor_Area_R | Pallidum_L | .980 |
| Fusiform_R | ACC_sup_L | .980 |
| Postcentral_R | ACC_sup_L | .980 |
| Occipital_Sup_L | SN_pc_R | .980 |
| Paracentral_Lobule_R | Putamen_L | .980 |
| Rolandic_Oper_L | Putamen_L | .979 |
| Pallidum_L | Temporal_Sup_R | .979 |
| Supp_Motor_Area_L | Putamen_L | .979 |
| OFCmed_L | Red_N_R | .979 |
| Putamen_L | Temporal_Inf_R | .979 |
| Parietal_Sup_R | Pallidum_L | .977 |
| N_Acc_R | Raphe_D | .977 |
| Occipital_Inf_R | SN_pr_R | .976 |
| Parietal_Sup_R | N_Acc_R | .976 |
| OFCant_R | Parietal_Inf_R | .975 |
| Insula_L | ACC_sup_L | .975 |
| Frontal_Mid_2_L | SupraMarginal_L | .975 |
| ParaHippocampal_R | Pallidum_L | .975 |
| Cingulate_Mid_L | Putamen_L | .975 |
| Frontal_Sup_Medial_L | Calcarine_L | .975 |
| Frontal_Sup_Medial_R | OFClat_R | .975 |
| OFCant_R | SupraMarginal_R | .975 |
| Pallidum_L | Red_N_R | .974 |
| Lingual_L | Pallidum_L | .974 |
| Olfactory_R | Cingulate_Post_R | .974 |
| ACC_sub_R | SN_pr_R | .974 |
| Frontal_Sup_2_L | Lingual_R | .974 |
| Precentral_L | N_Acc_R | .973 |
| N_Acc_L | Red_N_R | .973 |
| Cingulate_Mid_R | Pallidum_L | .972 |
| Amygdala_R | Red_N_R | .972 |
| Occipital_Inf_R | Putamen_L | .972 |
| Hippocampus_R | N_Acc_L | .972 |
| Fusiform_L | Putamen_L | .972 |
| Frontal_Inf_Orb_2_R | OFCant_R | .971 |
| SupraMarginal_L | Pallidum_L | .971 |
| Lingual_R | ACC_sub_R | .971 |
| Insula_R | SupraMarginal_R | .971 |
| Putamen_L | Temporal_Pole_Sup_L | .971 |
| Frontal_Mid_2_L | Calcarine_R | .970 |

| **Supplementary Table 11.** Partial correlations between BIS-11 total scores and nodal strength of regions within the relevant network, identified by NBS-Prediction, in P+ (P-CSO- > P+CSO+; weight threshold ≥ .97). | | | |
| --- | --- | --- | --- |
| **AAL Region** | ***r*** | ***p*** | ***p_FDR_*** |
| Pallidum_L | .190 | .267 | .507 |
| Putamen_L | .420 | .011 | .131 |
| Calcarine_R | .194 | .257 | .507 |
| ACC_pre_L | .187 | .274 | .507 |
| ACC_sup_L | .356 | .033 | .243 |
| OFCant_R | .163 | .341 | .532 |
| Lingual_R | .167 | .329 | .532 |
| Frontal_Mid_2_L | .254 | .135 | .354 |
| Precentral_L | .351 | .036 | .243 |
| SN_pr_R | .060 | .726 | .778 |
| Red_N_R | .246 | .148 | .357 |
| N_Acc_R | .344 | .040 | .243 |
| Occipital_Mid_L | .178 | .298 | .532 |
| SupraMarginal_L | .250 | .141 | .357 |
| Hippocampus_R | -.096 | .580 | .679 |
| Occipital_Inf_R | .316 | .060 | .258 |
| Fusiform_L | .264 | .119 | .354 |
| Calcarine_L | .255 | .133 | .354 |
| Frontal_Med_Orb_L | .085 | .620 | .708 |
| Frontal_Sup_Medial_L | .156 | .364 | .542 |
| SupraMarginal_R | .189 | .268 | .507 |
| Temporal_Mid_R | .272 | .108 | .354 |
| Frontal_Sup_Medial_R | .130 | .449 | .615 |
| Frontal_Sup_2_L | .111 | .519 | .632 |
| **Precentral_R** | **.571** | **<.001** | **.024** |
| ACC_pre_R | .187 | .274 | .507 |
| ACC_sub_R | .123 | .473 | .622 |
| Parietal_Sup_R | .415 | .012 | .131 |
| OFClat_R | .304 | .071 | .264 |
| Frontal_Sup_2_R | .113 | .511 | .631 |
| Frontal_Inf_Oper_R | .165 | .336 | .532 |
| ACC_sub_L | -.057 | .743 | .778 |
| Temporal_Pole_Sup_R | .393 | .018 | .158 |
| SN_pc_R | .121 | .481 | .622 |
| ACC_sup_R | .246 | .148 | .357 |
| Angular_R | .190 | .266 | .507 |
| Supp_Motor_Area_L | .268 | .114 | .354 |
| Frontal_Inf_Orb_2_R | .312 | .064 | .258 |
| Supp_Motor_Area_R | .338 | .044 | .243 |
| Olfactory_R | .078 | .653 | .726 |
| Frontal_Med_Orb_R | -.089 | .606 | .701 |
| Parietal_Sup_L | .326 | .052 | .250 |
| ParaHippocampal_R | .254 | .135 | .354 |
| Cingulate_Mid_L | .309 | .067 | .258 |
| Lingual_L | .261 | .125 | .354 |
| Postcentral_R | .483 | .003 | .125 |
| Caudate_L | .296 | .080 | .285 |
| Thal_PuL_R | .239 | .161 | .368 |
| Angular_L | .153 | .372 | .542 |
| Frontal_Inf_Oper_L | .061 | .722 | .778 |
| Frontal_Mid_2_R | .141 | .413 | .582 |
| Rolandic_Oper_R | .433 | .008 | .131 |
| N_Acc_L | .041 | .811 | .829 |
| Raphe_D | .119 | .489 | .622 |
| Temporal_Sup_R | .233 | .171 | .381 |
| Fusiform_R | .334 | .046 | .243 |
| Frontal_Inf_Tri_R | .150 | .384 | .551 |
| Postcentral_L | .431 | .009 | .131 |
| Occipital_Mid_R | .313 | .063 | .258 |
| Cingulate_Post_L | .055 | .751 | .778 |
| Parietal_Inf_R | .177 | .300 | .532 |
| Insula_L | .242 | .155 | .362 |
| Rolandic_Oper_L | .401 | .015 | .153 |
| Frontal_Inf_Tri_L | .100 | .562 | .667 |
| Frontal_Inf_Orb_2_L | .258 | .129 | .354 |
| OFCant_L | .125 | .466 | .622 |
| OFCmed_R | .165 | .337 | .532 |
| OFCmed_L | .190 | .266 | .507 |
| Rectus_R | -.121 | .484 | .622 |
| OFCpost_R | .191 | .266 | .507 |
| Insula_R | .161 | .347 | .533 |
| Hippocampus_L | -.034 | .846 | .855 |
| Amygdala_R | .277 | .102 | .350 |
| Parietal_Inf_L | .139 | .418 | .582 |
| Occipital_Inf_L | .078 | .651 | .726 |
| Occipital_Sup_L | .107 | .535 | .644 |
| OFClat_L | .020 | .907 | .907 |
| Cingulate_Mid_R | .155 | .367 | .542 |
| Cingulate_Post_R | .058 | .738 | .778 |
| Paracentral_Lobule_R | .424 | .010 | .131 |
| Precuneus_R | .171 | .318 | .532 |
| Temporal_Mid_L | .069 | .690 | .758 |
| Temporal_Pole_Sup_L | .360 | .031 | .243 |
| Putamen_R | .427 | .009 | .131 |
| Caudate_R | .325 | .053 | .250 |
| Precuneus_L | .165 | .337 | .532 |
| Temporal_Inf_R | .339 | .043 | .243 |
| Thal_PuM_L | .171 | .317 | .532 |
| Raphe_M | .116 | .501 | .628 |
| **Note:** Dummy-coded site variables were corrected in partial correlation. FDR correction was applied across brain regions within each contrast. Significant correlations are highlighted. | | | |

### **Supplementary References**

1. Fornito A, Zalesky A, Bullmore ET (2016): *Fundamentals of Brain Network Analysis*. Elsevier. https://doi.org/10.1016/C2012-0-06036-X

2. Preuss UW, Rujescu D, Giegling I, Watzke S, Koller G, Zetzsche T, *et al.* (2008): Psychometrische Evaluation der deutschsprachigen Version der Barratt-Impulsiveness-Skala. *Nervenarzt* 79: 305–319.

3. Patton JH, Stanford MS, Barratt ES (1995): Factor structure of the Barratt impulsiveness scale. *J Clin Psychol* 51: 768–774.

4. Rolls ET, Huang C-C, Lin C-P, Feng J, Joliot M (2020): Automated anatomical labelling atlas 3. *NeuroImage* 206: 116189.

5. Yeo BTT, Krienen FM, Sepulcre J, Sabuncu MR, Lashkari D, Hollinshead M, *et al.* (2011): The organization of the human cerebral cortex estimated by intrinsic functional connectivity. *J Neurophysiol* 106: 1125–1165.
